# Insights from a mobile healthcare clinic in Madagascar: Diversity and dynamics of viruses from febrile individuals in rural subsistence communities

**DOI:** 10.64898/2026.10.07.757397

**Authors:** Elise N. Paietta, Andriantiana Tsirimanana, Rachel A. Johnston, Tanguy F. Razafiarimihoby, Olesya Ravaoharimbola, Simona Kraberger, Camille M.M. DeSisto, Emmanuel Silva Orozco, Melodi Anahtar, Brian McAdoo, Eric Baitchman, Sakib Burza, Matthew Scotch, Arvind Varsani

## Abstract

Communities living in biodiversity hotspots are at increased risk of pathogen exposure and spread due to insufficient access to nutrition, clean water, and sanitation. Viruses, in particular, have high epidemic and spillover potential, and widespread health impacts, yet, viral diversity and dynamics remain understudied in many high-risk regions. In Madagascar, a hotspot with unparalleled biodiversity, socioeconomic constraints, and a heavy burden of infectious diseases, ongoing healthcare initiatives provide opportunities to survey circulating viral diversity. Here, we used a metagenomic workflow to characterize the viromes of oral/nasal swabs from febrile individuals visiting a mobile healthcare clinic in southeastern Madagascar, of which 49% were malaria-positive (*Plasmodium falciparum*). We characterized sequences of human-infecting viruses from twelve families *(Adenoviridae, Anelloviridae, Orthoherpesviridae, Papillomaviridae, Polyomaviridae*, *Astroviridae, Coronaviridae, Matonaviridae, Orthomyxoviridae, Paramyxoviridae, Picornaviridae,* and *Sedoreoviridae*). These viral sequences generally showed high similarity to known lineages circulating globally, but multiple viruses considered rare or emerging were identified (e.g. enterovirus C105, enterovirus C117). While we did not detect a significant association between malaria status and pathogenic virus presence, younger individuals had higher odds of having a pathogenic virus. Nonpathogenic viruses in *Anelloviridae* were prevalent and diverse with richness ranging from 0 to 70. Individuals with *P. falciparum* or a pathogenic virus tended to share more anelloviruses. We also detected viruses that are protist-associated, plant-infecting, or with unknown hosts. Overall, our work documents extensive viral diversity in Madagascar, highlighting how metagenomic workflows can be used to build viral genomic resources for pathogen surveillance in regions with high infectious disease burden.

## Introduction

Lack of access to healthcare, adequate nutrition, clean water, and sanitation can increase pathogen exposure, prolong infection duration, facilitate pathogen spread, and result in more severe clinical manifestations of disease (Bryce et al., 2005; Freeman et al., 2017; Prüss-Ustün et al., 2014). Many pathogenic viruses are of particular concern for communities without sufficient resource access due to their high transmissibility (e.g., measles virus), rapid evolution (e.g., SARS-CoV-2), impacts on host immune systems (e.g., HIV), and potential to establish persistent infections (e.g., herpesviruses). Further, factors such as malnutrition and anemia (Osman, 2016; Rice et al., 2000; Stewart et al., 2020) and co-infections (e.g., co-infection of *Plasmodium* and Epstein-Barr virus increases the risk of developing Burkitt lymphoma (Mawson and Majumdar, 2017)) can contribute to immunosuppression, exacerbating the progression and severity of many viral infections. Despite the interactions between socioeconomic conditions and viral exposure, little is known about viral dynamics in many high-poverty regions due to insufficient access to resources for simultaneously tracking diverse viruses of interest.

Madagascar is an island nation off Africa’s eastern coast, home to an estimated 33 million people. Madagascar is recognized for its evolutionarily unique biodiversity with ∼90% endemism of plants and animals. Malagasy people additionally have a distinct, rich history with both Asian and African ancestors, an Austronesian language, and vibrant cultures influenced by environmental variability across the island (Allibert, 2008; Dewar and Richard, 2012).

Madagascar is also burdened by extreme poverty and has been ranked 120^th^ out of 123 countries in the Global Hunger Index (2025). According to the World Health Organization (2021), behind strokes, respiratory tract infections and diarrheal diseases are the top causes of death for people in Madagascar. Studies in Madagascar have found that the majority of respiratory tract infections are due to infection (and frequently co-infection) from diverse viruses, including adenoviruses, coronaviruses, influenza viruses, respiratory syncytial virus, and rhinoviruses (Hoffmann et al., 2012; Lockwood et al., 2026; Rajatonirina et al., 2013b; Randriambolamanantsoa et al., 2024; Razanajatovo et al., 2018; Razanajatovo et al., 2022; Razanajatovo et al., 2011). While intestinal parasites and bacteria are additionally major etiological agents of diarrheal disease and thus drivers of chronic anemia and malnutrition in people (Howard et al., 2007; Prendergast and Kelly, 2016; Randremanana et al., 2012), rotaviruses, noroviruses, adenoviruses, and astroviruses have also been recognized as common causes of diarrheal disease (Randremanana et al., 2016; Razanajatovo et al., 2023; Zohdy et al., 2015). Oral and nasal swab samples are key for detecting respiratory viruses (Cui et al., 2024; Graf et al., 2016); enteric viruses can also be detected, though usually at lower rates than from stool samples (Anfruns-Estrada et al., 2020; Zhuo et al., 2018).

Despite the role of viruses in human morbidity and mortality in Madagascar, genomic resources for viruses characterized in Malagasy people are relatively limited with 45 complete and 105 near-complete virus genomes out of 1,984 nucleotide virus sequence records currently on NCBI Virus (Brister et al., 2015). Recent studies have particularly expanded genetic resources for respiratory syncytial virus, measles virus, and rubella virus in Madagascar (Lockwood et al., 2026; Razafindratsimandresy et al., 2026). For reference, countries with a similar population size to Madagascar have least an order of magnitude more virus genetic resources deposited (NCBI Virus), including Cameroon (26,772 nucleotide virus sequence records), Malaysia (12,334), and Peru (50,949). With only ∼3.3% of the Malagasy government’s GDP reported to be allocated to health (2022), ongoing humanitarian aid plays a substantial role in meeting basic health needs in many regions. As organizations and communities across Madagascar unite to improve healthcare access and infrastructure, these healthcare initiatives provide an opportunity to survey the landscape of viruses impacting high-poverty remote communities and to build better genomic resources for Madagascar.

As part of an initiative to broaden our understanding of viral diversity in Madagascar, we worked with the non-governmental organization Health In Harmony, to initiate a study whereby individuals visiting a mobile healthcare clinic in southeastern Madagascar and presenting with febrile illness were invited to participate in the study and provide oral and nasal swab samples. The sampled communities are living within and around the Manombo Special Reserve, a biodiverse yet rapidly degrading protected area home to endemic and non-native mammals including non-human primates, rodents, bats and carnivores. Deforestation across the island has promoted favorable conditions for vectors such as mosquitoes, and many regions of Madagascar have high malaria burdens, with many households (>50%) in southeastern Madagascar found to have multiple infected individuals (Rice et al., 2021). However, increased use of malaria rapid diagnostic testing has revealed a higher proportion of febrile patients that are *Plasmodium*-negative, and studies in Madagascar have begun to identify numerous viral pathogens present in these individuals using targeted panels (e.g. Rift Valley fever virus, Varicella Zoster virus, respiratory syncytial virus, influenza viruses A and B) (Guillebaud et al., 2018).

Using viral metagenomic approaches, our work capitalizes on recent global efforts to build a more comprehensive understanding of the human virome through high-quality genomic characterization of both pathogenic and non-pathogenic human-infecting viruses. While many pathogenic viruses have direct effects on host immune systems, non-pathogenic viruses additionally influence host immunity and some are considered commensal. Here, we examine both RNA and DNA viral diversity in oral and nasal swabs, as well as the factors shaping viral infection and co-occurrence dynamics in remote Malagasy communities, while generating extensive genomic data in an understudied, biodiverse region. This project assesses the similarity of viruses in communities in Madagascar to those found globally circulating in human populations, while additionally demonstrating the strength of partnerships with healthcare clinics to perform viral surveillance in rural, resource-limited settings.

## Methods and materials

### Sample collection

The human sampling described here was approved through the Comité Malgache d’Éthique pour les Sciences et les Technologies (CMEST) under permit N°194-2024 /AM/CMEST-P/ in Madagascar, the Duke University Campus Institutional Review Board under protocol #2025-0019, and the Arizona State University Institutional Review Board under study ID #STUDY00022785. The human sampling was conducted through a partnership with a mobile healthcare clinic of non-governmental organization Health In Harmony which serves 31 villages (12 mobile clinic sites) within and surrounding the Manombo Special Reserve in the Farafangana region of southeastern Madagascar. This mobile clinic has been working with villages since 2020. Those seeking care often present with febrile illness symptoms, which, given high malaria prevalence in the region, warrants a malaria rapid diagnostic test (RDT) (Meriscreen One Step test for Malaria Pf/Pan Ag; Meril Diagnostics, India). This RDT is an immunochromatographic assay for qualitative *Plasmodium* infection by *Plasmodium falciparum* (HRP2 antigens) and *P. falciparum, P. vivax, P. malariae,* and *P. ovale* (pan-pLDH antigens).

In September 2024, oral and nasal swabs and associated de-identified metadata (village, age, gender, malaria status) were collected from the first 20 consenting individuals presenting with a fever, at times accompanied by weakness, headaches, or muscle aches at each of the 12 clinic sites (n = 240 samples). Sampling occurred independently of RDT results. The oral and nasal swabs were collected using sterile flocked swabs (Puritan Medical Products, USA) and stored in 1 ml of DNA/RNA Shield (Zymo, USA). An oral swab and nasal swab from the same individual were stored together in one cryotube as one sample for ease of transport. All metadata were unavailable for 14 individuals and malaria status results, specifically, were unavailable for 70 individuals due to the fast pace of the mobile clinic environment. Samples were transported in liquid nitrogen and subsequently maintained in a −80°C freezer until nucleic acid extraction.

### Nucleic acid extraction and sequencing

After brief gentle vortexing, DNA was extracted from 200 µl of each sample with the Roche High Pure Viral Nucleic Acid Kit (Roche Diagnostics, USA) while RNA was extracted from 200 µl of each sample with the RNeasy Plus Mini Kit (Qiagen, Germany). DNA extracts were amplified using the Templiphi rolling circle amplification kit (Cytiva Life Sciences, USA) to enrich for circular DNA viruses. For DNA sequencing library preparation, rolling circle-amplified and non-amplified DNA extracts were combined (50:50) and used as input to generate Illumina sequencing libraries (2×150bp) using the Illumina DNA Prep (M) Tagmentation Kit (Illumina, USA). For RNA, RNA extract was used as input to generate Illumina sequencing libraries (2×150bp) using the Illumina Stranded Total RNA Prep with Ligation, Ribo-Zero Plus Microbiome Kit (Illumina, USA). Libraries were sequenced on the NovaSeq X Plus with Duke Sequencing and Genomic Technologies (Duke University, USA) or Psomagen Inc. (USA).

Reads were trimmed with Trimmomatic-0.39 (Bolger et al., 2014) prior to *de novo* assembly using MEGAHITv.1.2.9 (Li et al., 2015). Terminal redundancy was used to identify circular contigs. Contigs were analyzed for viral-like sequences with Diamond (Buchfink et al., 2015) BLASTx against a local viral RefSeq database (release 230). Viral sequences were annotated with CenoteTaker3 (Tisza et al., 2021; Tisza et al., 2025; Tisza et al., 2026) in addition to manual curation using available human-infecting virus genomes as references.

### Phylogenetic and sequence similarity analyses

For viruses selected for further analysis, datasets of nucleotide sequences were assembled together with sequences of viruses identified in this study and aligned with MAFFT v7 (Katoh and Standley, 2013). Reference sequences from NCBI were downloaded for each virus group of interest except for influenza C virus where sequences were downloaded from GISAID (Shu and McCauley, 2017). Maximum likelihood phylogenetic trees were built for most viruses using IQ-Tree 2 (Minh et al., 2020) with the ModelFinder (Kalyaanamoorthy et al., 2017) option and approximate likelihood ratio test (aLRT; 1,000 replicates) branch support. Best-fit substitution models were determined for the rubella virus dataset (whole genome: TN+F+I+G4; *E1* nt: TN+F+G4), enterovirus C105 and C117 datasets (*VP1* nt: TIM2e+I+G4), Madahom bibyvirus 1 datset (polyprotein aa: LG+F+I+G4), human astrovirus dataset (*orf2* nt: TIM2+F+G4), rotavirus A dataset (*VP4* nt: TVM+F+I+G4), human parainfluenza virus (HPIV)-2 dataset (whole genome: GTR+F+R2), HPIV-3 dataset (whole genome: TIM+F+R2), and redondovirus dataset (Rep aa: LG+F+I+G4). All phylogenetic trees were rooted with representative outgroup sequences (specified in figure captions). Branches with <0.8 aLRT support were collapsed with TreeGraph2 (Stöver and Müller, 2010) and trees were visualized with iToL (Letunic and Bork, 2021). Rotavirus A genotyping was confirmed with the RIVM genotyping tool Version 0.1. For HPIV-2 and −3, genotype labelling was based on Zhu et al. (Zhu et al., 2024).

For reconstructing the evolutionary history of influenza C virus we used BEAST v1.10.5 (Baele et al., 2025) to perform Bayesian phylogenetic inference under a HKY + [\GAMMA] model of DNA substitution, an uncorrelated relaxed molecular clock (Drummond et al., 2006), and a Bayesian Skygrid population growth prior (Hill and Baele, 2019). We recorded two independent Markov chain Monte Carlo (MCMC) simulations that had a chain length of 5×10^8^ and a sampling frequency of 5×10^4^. We used Tracer v1.7.2 (Rambaut et al., 2018) to check for model convergence including that all ESS parameters were ≥ 200 and compare the independent runs. We combined posterior log and tree files via LogCombiner v1.10.5 (Baele et al., 2025) and created a maximum clade credibility (MCC) tree with TreeAnnotator v1.10.5 (Baele et al., 2025). Additionally, we used Tracer v1.7.2 to produce a plot of the effective population size of influenza C viruses under a Bayesian Skygrid demographic prior. We used the Baltic library in Python for visualization of the MCC tree. Sequence similarity was analyzed with SDT v1.3 (Sequence Demarcation Tool) (Muhire et al., 2014).

### Distribution of virus genomes across samples

Virus genomes were clustered into virus operational taxonomic units (vOTUs) using CD-Hit (Fu et al., 2012) with a 95% genome-wide average nucleotide identity threshold (Roux et al., 2019). Illumina sequencing raw reads from all samples were then mapped to a representative genome from each vOTU using CoverM (Aroney et al., 2025) (with -min-read-percent-identity 95%). This output was used to determine the presence or absence of each vOTU across all samples using a cut-off of ≥75% genome coverage as a proxy of presence. A matrix of vOTU presence absence for both RNA and DNA viruses across all samples was used as a basis for alpha- and beta-diversity metrics and network analyses. Regression modeling using pathogenic virus presence was computed using the R package *lme4* (Bates et al., 2015).

### Anellovirus co-occurrence analyses

To investigate the predictors of anellovirus vOTU co-occurrence (i.e., sharing) among individuals, we created a weighted anellovirus co-occurrence network, where nodes represented individuals and weighted edges represented the number of anellovirus vOTUs they had in common. Next, we fitted a valued ERGM (Exponential Random Graph Model) with a Poisson reference using the R package *ergm* (Hunter et al., 2008; Krivitsky, 2012) to test the predictive effects of pathogen co-occurrence, demographic traits, and spatial proximity on the number of anellovirus vOTUs shared between pairs of individuals. ERGMs are generative models of network attributes and structure which implement Markov chain Monte Carlo maximum likelihood parameter estimation. We included a suite of terms in the model, including binary node factor terms for malaria positivity and pathogenic virus infection. An individual was considered to have a pathogenic virus infection if they were carrying at least one pathogenic virus (DNA or RNA virus). Age was included both as a node covariate and through an “absdiff” term which measures the difference in age between two nodes. Similarly, sex was included as a node factor and using a “nodematch” term which accounts for sex-based homophily (i.e., whether pairs of individuals are likely to share more anelloviruses if they are the same sex). Spatial proximity was accounted for in an edge covariate term for the geographic distance between resident villages of the individuals, which we calculated using GPS coordinates and the R package *geosphere* (Hijmans et al., 2017). We also included network structural terms: “sum” (i.e., the total number of edges, similar to the “intercept”) and “nonzero” (i.e., the number of pairs of nodes that share any edge). Note that all terms included are dyad-independent; we estimated the associations between covariates and virus co-occurrence without modelling network structure beyond the “sum” and “nonzero” terms. In the main model, we excluded individuals with missing covariate data, for a total of 170 individuals. Due to limited sampling of malaria positivity, we also fitted an additional model without malaria and pathogenic virus covariates, for a total of 226 people (Supplementary Figure 3).

## Results and discussion

### Sampling overview

As part of a broader One Health study (Paietta et al., 2026a; Paietta et al., 2026b; Paietta et al., 2026c), oral/nasal swabs were collected from individuals (n = 240) presenting with febrile illness at a mobile healthcare clinic in southeastern Madagascar in September 2024 with demographic metadata available for 226 individuals. Twenty samples were collected per mobile clinic site, representing individuals from 29 out of 31 villages in the region (Figure 1). For individuals with reported results, 49.4% had a positive malaria status via RDT with tests indicating infection with either ‘*Plasmodium falciparum*’ or ‘*Plasmodium falciparum* and other’ (Figure 1A, Supplementary Figure 1A). Malaria prevalence and pathogenic virus detection frequency varied widely between villages (0-100%), despite samples being collected within a 20-day timespan (Figure 1A, B). Individuals in this study ranged in age from seven months to 88 years, with many samples coming from individuals aged <10 years (*n* = 87) (Supplementary Figure 1B). Additional samples were distributed across the following age groups: 10–20 years (*n* = 35), 20–30 years (*n* = 38), 30–40 years (*n* = 22), 40–50 years (*n* = 19), 50–60 years (*n* = 13), 60–70 years (*n* = 6), 70–80 years (*n* = 5), and 80–90 years (*n* = 1) (Supplementary Figure 1B). The dataset is skewed towards females (*n* = 155) in comparison with males (*n* = 71), reflecting gender differences in mobile clinic usage, particularly among adults ≥20 years old (n_female_ = 75, n_male_ = 29) as women frequently bring children to the clinic and are thus more likely to seek clinic services for themselves (Supplementary Figure 1B). Geographically, these samples span communities living at the edges of biodiverse forests (the Manombo Special Reserve) including more inland lowland rainforest and more coastal littoral forest regions (Figure 1). We emphasize that viruses in these samples primarily represent oral and nasal cavity-associated viruses and are thus a subset of the viruses circulating in these rural communities.

**Figure 1.**
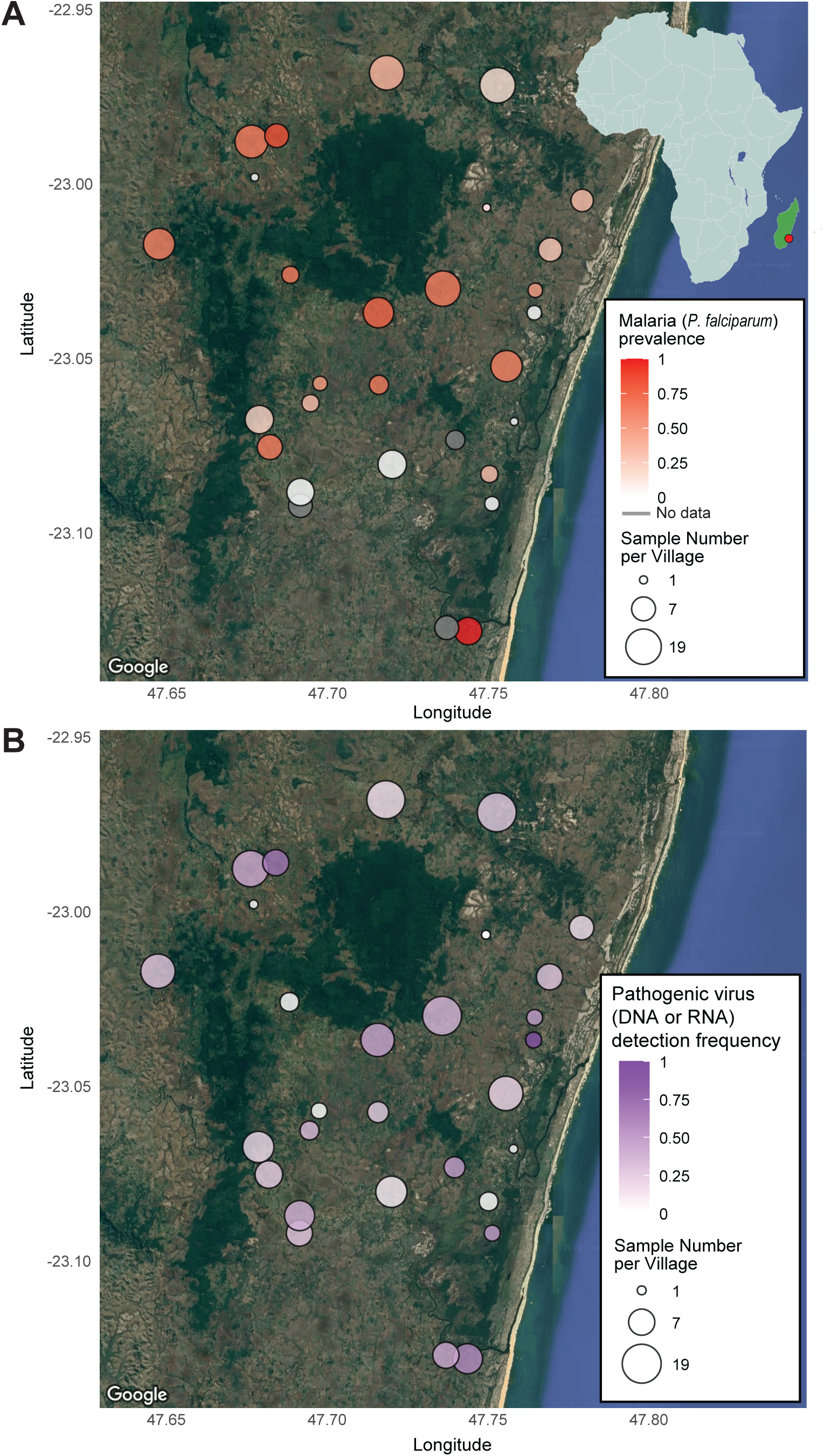
**(A)** Map depicting village locations with points sized by sample number and colored by malaria prevalence (white = 0% prevalence, red = 100% prevalence, gray = no data available). The upper right corner depicts Madagascar’s location in relation to mainland Africa with a red point highlighting the sampling location in southeastern Madagascar. **(B)** Map depicting village locations with points sized by sample number and colored by pathogenic virus detection frequency (i.e., the proportion of people in that village with at least one pathogenic virus vOTU present) (white = 0% prevalence, purple = 100% prevalence).

Viral metagenomic analysis of oral/nasal swab DNA and RNA resulted in the identification of viruses across five DNA viral families with human-infecting members: *Adenoviridae* (n = 2 coding-complete sequences)*, Anelloviridae* (n = 530)*, Orthoherpesviridae* (n = 2)*, Papillomaviridae* (n = 5), and *Polyomaviridae* (n = 9) (Table 1), and seven RNA viral families with human-infecting members: *Astroviridae* (n = 1)*, Coronaviridae* (n = 4)*, Matonaviridae* (n = 2)*, Orthomyxoviridae* (n = 2, 7 segments each)*, Paramyxoviridae* (n = 8)*, Picornaviridae* (n = 22), and *Sedoreoviridae* (n = 1, 11 segments) (Table 2; Supplementary Table 1). Via oral swabs samples, we also identified other eukaryote-infecting viruses, including protist-infecting viruses in the *Redondoviridae* family (n =37), plant-infecting viruses in the *Alphaflexviridae* (n = 1), *Geminiviridae* (n = 7), and *Virgaviridae* (n = 14) families, along with restvirus (statovirus-like) (n = 2) and tombusvirus-like (n = 1) sequences with unknown hosts (Table 3; Supplementary Table 1). Thus, we characterized coding-complete sequences of dsDNA, ssDNA, dsRNA, ssRNA(-), and ssRNA(+) viruses. The majority of DNA virus sequences are members of the *Anelloviridae* family and represent 446 vOTUs. The other DNA virus sequences represent 31 vOTUs across five families. The RNA virus sequences represent 24 vOTUs across seven families and two non-established taxa. Mean coverage for DNA viruses ranged from 4.3 to 120612.8 (Supplementary Table 1). Mean coverage for RNA viruses ranged from 6.9 to 16641.4 (Supplementary Table 1). Viruses were detected in the oral/nasal swabs of 179 out of 240 individuals, likely due to the low biomass obtained from swabbing.

**Table 1.** Summary of human-infecting DNA viruses identified in oral/nasal swab samples and their similarity to published sequences.

| Virus Family | Virus Species | Virus Name | Accession | Length (nt) | BLASTn highest similarity hit | Percent identity (%) | Query cover (%) |
| --- | --- | --- | --- | --- | --- | --- | --- |
| **Polyomaviridae** | *Alphapolyomavirus quintihominis* | Merkel cell PyV | **PZ726174** | 5392 | ON023014 | 99.85 | 100 |
|  |  |  | **PZ726173** | 5392 | ON023013 | 99.74 | 100 |
|  |  |  | **PZ726172** | 5382 | ON023014 | 99.07 | 100 |
| **Polyomaviridae** | *Betapolyomavirus quartihominis* | WU PyV | **PZ726221** | 5229 | GU296367 | 100 | 100 |
|  |  |  | **PZ726220** | 5229 | GU296367 | 100 | 100 |
|  |  |  | **PZ726219** | 5229 | GU296367 | 100 | 100 |
|  |  |  | **PZ726218** | 5229 | GU296367 | 100 | 100 |
|  |  |  | **PZ726217** | 5228 | PQ335160 | 98.83 | 100 |
| **Polyomaviridae** | *Deltapolyomavirus undecihominis* | STL PyV | **PZ726216** | 4776 | JX463183 | 98.01 | 100 |
| **Papillomaviridae** | *Betapapillomavirus 1* | HPV type 47 | **PZ726168** | 7723 | M32305 | 91.59 | 100 |
| **Papillomaviridae** | *Betapapillomavirus 3* | HPV type 49 | **PZ726169** | 7575 | X74480 | 98.75 | 100 |
| **Papillomaviridae** | *Gammapapillomavirus* sp. | HPV-mSK_227 | **PZ726171** | 7263 | MH777366 | 97.51 | 100 |
| **Papillomaviridae** | *Gammapapillomavirus* sp. | HPV-mSK_020 | **PZ726170** | 7253 | MH777168 | 96.61 | 100 |
| **Papillomaviridae** | *Betapapillomavirus 2* | HPV type 23 | **PZ726167** | 7324 | U31781 | 99.19 | 100 |
| **Adenoviridae** | *Mastadenovirus caesari* | Human adenovirus 89 | **PZ726818** | 35906 | OR753116 | 99.66 | 100 |
| **Adenoviridae** | *Mastadenovirus caesari* | Human adenovirus 1 | **PZ726819** | 35986 | OQ518269 | 99.53 | 100 |
| **Orthoherpesviridae** | *Cytomegalovirus humanbeta5* | Human betaherpesvirus 5 (HCMV) | **PZ726821** | 234144 | KP745634 | 98.02 | 100 |
| **Orthoherpesviridae** | *Lymphocryptovirus humangamma4* | Human gammaherpesvirus 4 (EBV) | **PZ726820** | 167614 | KX125053 | 99.11 | 100 |

**Table 2.**
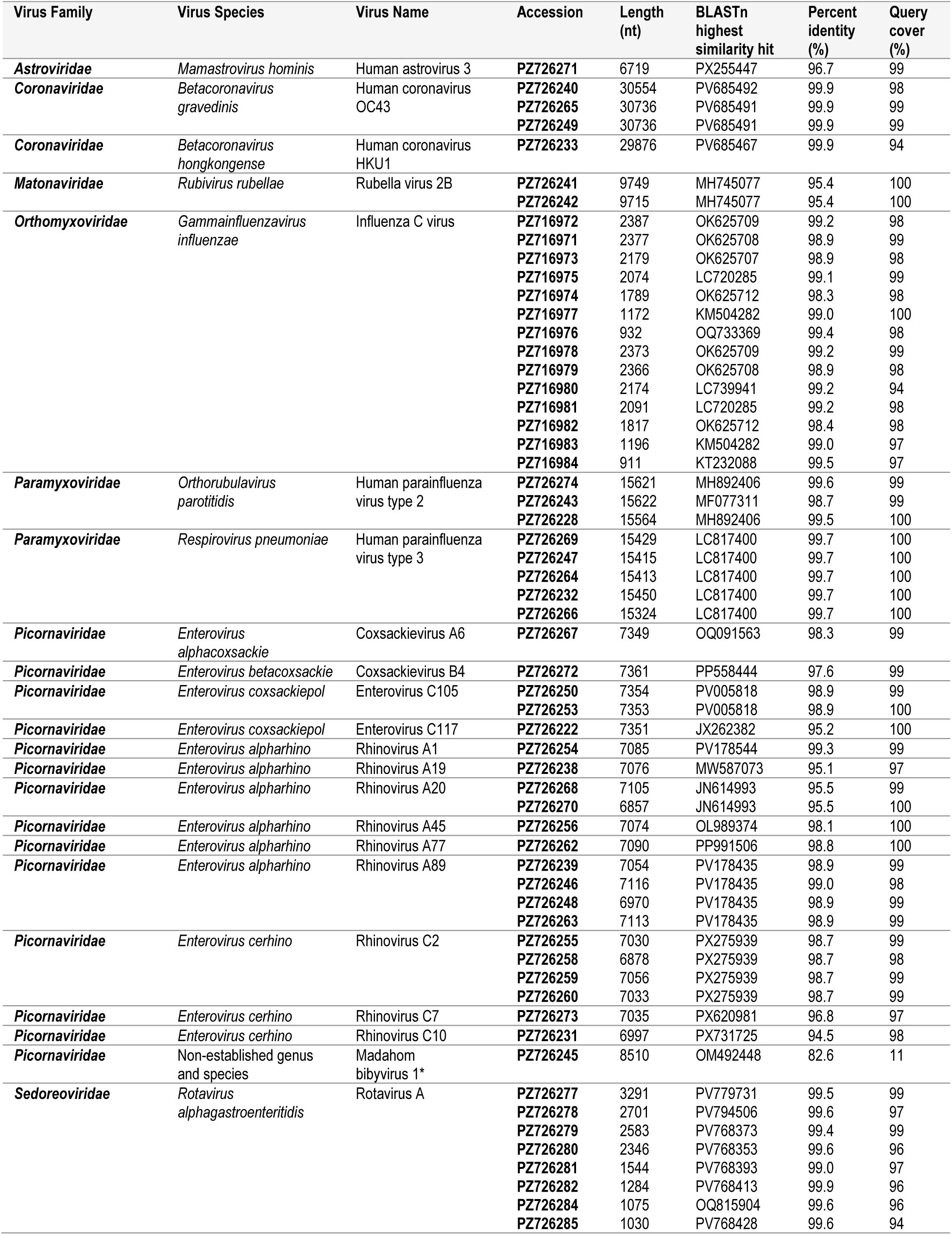
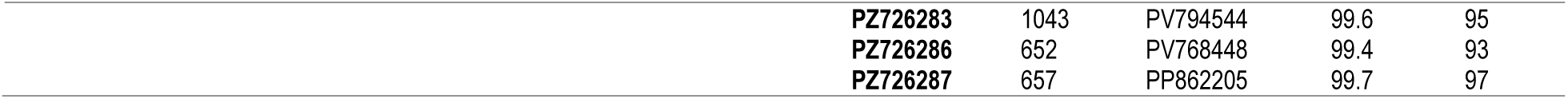
Summary of human-infecting RNA viruses identified in oral/nasal swab samples and their similarity to published sequences. *Putative mammalian host, unconfirmed.

| Virus Family | Virus Species | Virus Name | Accession | Length (nt) | BLASTn highest similarity hit | Percent identity (%) | Query cover (%) |
| --- | --- | --- | --- | --- | --- | --- | --- |
| **Astroviridae** | *Mamastrovirus hominis* | Human astrovirus 3 | PZ726271 | 6719 | PX255447 | 96.7 | 99 |
| **Coronaviridae** | *Betacoronavirus gruedinis* | Human coronavirus OC43 | PZ726240 | 30554 | PV685492 | 99.9 | 98 |
|  |  |  | PZ726265 | 30736 | PV685491 | 99.9 | 99 |
|  |  |  | PZ726249 | 30736 | PV685491 | 99.9 | 99 |
| **Coronaviridae** | *Betacoronavirus hongkongense* | Human coronavirus HKU1 | PZ726233 | 29876 | PV685467 | 99.9 | 94 |
| **Matonaviridae** | *Rubivirus rubellae* | Rubella virus 2B | PZ726241 | 9749 | MH745077 | 95.4 | 100 |
|  |  |  | PZ726242 | 9715 | MH745077 | 95.4 | 100 |
| **Orthomyxoviridae** | *Gammmainfluenzavirus influenzae* | Influenza C virus | PZ716972 | 2387 | OK625709 | 99.2 | 98 |
|  |  |  | PZ716971 | 2377 | OK625708 | 98.9 | 99 |
|  |  |  | PZ716973 | 2179 | OK625707 | 98.9 | 98 |
|  |  |  | PZ716975 | 2074 | LC720285 | 99.1 | 99 |
|  |  |  | PZ716974 | 1789 | OK625712 | 98.3 | 98 |
|  |  |  | PZ716977 | 1172 | KM504282 | 99.0 | 100 |
|  |  |  | PZ716976 | 932 | OQ733369 | 99.4 | 98 |
|  |  |  | PZ716978 | 2373 | OK625709 | 99.2 | 99 |
|  |  |  | PZ716979 | 2366 | OK625708 | 98.9 | 98 |
|  |  |  | PZ716980 | 2174 | LC739941 | 99.2 | 94 |
|  |  |  | PZ716981 | 2091 | LC720285 | 99.2 | 98 |
|  |  |  | PZ716982 | 1817 | OK625712 | 98.4 | 98 |
|  |  |  | PZ716983 | 1196 | KM504282 | 99.0 | 97 |
|  |  |  | PZ716984 | 911 | KT232088 | 99.5 | 97 |
| **Paramyxoviridae** | *Orthorubulavirus parotitidis* | Human parainfluenza virus type 2 | PZ726274 | 15621 | MH892406 | 99.6 | 99 |
|  |  |  | PZ726243 | 15622 | MF077311 | 98.7 | 99 |
|  |  |  | PZ726228 | 15564 | MH892406 | 99.5 | 100 |
| **Paramyxoviridae** | *Respirovirus pneumoniae* | Human parainfluenza virus type 3 | PZ726269 | 15429 | LC817400 | 99.7 | 100 |
|  |  |  | PZ726247 | 15415 | LC817400 | 99.7 | 100 |
|  |  |  | PZ726264 | 15413 | LC817400 | 99.7 | 100 |
|  |  |  | PZ726232 | 15450 | LC817400 | 99.7 | 100 |
|  |  |  | PZ726266 | 15324 | LC817400 | 99.7 | 100 |
| **Picornaviridae** | *Enterovirus alphacoxsackie* | Coxsackievirus A6 | PZ726267 | 7349 | OQ091563 | 98.3 | 99 |
| **Picornaviridae** | *Enterovirus betacoxsackie* | Coxsackievirus B4 | PZ726272 | 7361 | PP558444 | 97.6 | 99 |
| **Picornaviridae** | *Enterovirus coxsackiepol* | Enterovirus C105 | PZ726250 | 7354 | PV005818 | 98.9 | 99 |
|  |  |  | PZ726253 | 7353 | PV005818 | 98.9 | 100 |
| **Picornaviridae** | *Enterovirus coxsackiepol* | Enterovirus C117 | PZ726222 | 7351 | JX262382 | 95.2 | 100 |
| **Picornaviridae** | *Enterovirus alpharhino* | Rhinovirus A1 | PZ726254 | 7085 | PV178544 | 99.3 | 99 |
| **Picornaviridae** | *Enterovirus alpharhino* | Rhinovirus A19 | PZ726238 | 7076 | MW587073 | 95.1 | 97 |
| **Picornaviridae** | *Enterovirus alpharhino* | Rhinovirus A20 | PZ726268 | 7105 | JN614993 | 95.5 | 99 |
|  |  |  | PZ726270 | 6857 | JN614993 | 95.5 | 100 |
| **Picornaviridae** | *Enterovirus alpharhino* | Rhinovirus A45 | PZ726256 | 7074 | OL989374 | 98.1 | 100 |
| **Picornaviridae** | *Enterovirus alpharhino* | Rhinovirus A77 | PZ726262 | 7090 | PP991506 | 98.8 | 100 |
| **Picornaviridae** | *Enterovirus alpharhino* | Rhinovirus A89 | PZ726239 | 7054 | PV178435 | 98.9 | 99 |
|  |  |  | PZ726246 | 7116 | PV178435 | 99.0 | 98 |
|  |  |  | PZ726248 | 6970 | PV178435 | 98.9 | 99 |
|  |  |  | PZ726263 | 7113 | PV178435 | 98.9 | 99 |
| **Picornaviridae** | *Enterovirus cerhino* | Rhinovirus C2 | PZ726255 | 7030 | PX275939 | 98.7 | 99 |
|  |  |  | PZ726258 | 6878 | PX275939 | 98.7 | 98 |
|  |  |  | PZ726259 | 7056 | PX275939 | 98.7 | 99 |
|  |  |  | PZ726260 | 7033 | PX275939 | 98.7 | 99 |
| **Picornaviridae** | *Enterovirus cerhino* | Rhinovirus C7 | PZ726273 | 7035 | PX620981 | 96.8 | 97 |
| **Picornaviridae** | *Enterovirus cerhino* | Rhinovirus C10 | PZ726231 | 6997 | PX731725 | 94.5 | 98 |
| **Picornaviridae** | Non-established genus and species | Madahom bibyavirus 1* | PZ726245 | 8510 | OM492448 | 82.6 | 11 |
| **Sedoreoviridae** | *Rotavirus alphagastroenteritidis* | Rotavirus A | PZ726277 | 3291 | PV779731 | 99.5 | 99 |
|  |  |  | PZ726278 | 2701 | PV794506 | 99.6 | 97 |
|  |  |  | PZ726279 | 2583 | PV768373 | 99.4 | 99 |
|  |  |  | PZ726280 | 2346 | PV768353 | 99.6 | 96 |
|  |  |  | PZ726281 | 1544 | PV768393 | 99.0 | 97 |
|  |  |  | PZ726282 | 1284 | PV768413 | 99.9 | 96 |
|  |  |  | PZ726284 | 1075 | OQ815904 | 99.6 | 96 |
|  |  |  | PZ726285 | 1030 | PV768428 | 99.6 | 94 |

|  |  |  |  |  |
| --- | --- | --- | --- | --- |
| **PZ726283** | 1043 | PV794544 | 99.6 | 95 |
| **PZ726286** | 652 | PV768448 | 99.4 | 93 |
| **PZ726287** | 657 | PP862205 | 99.7 | 97 |

**Table 3.**
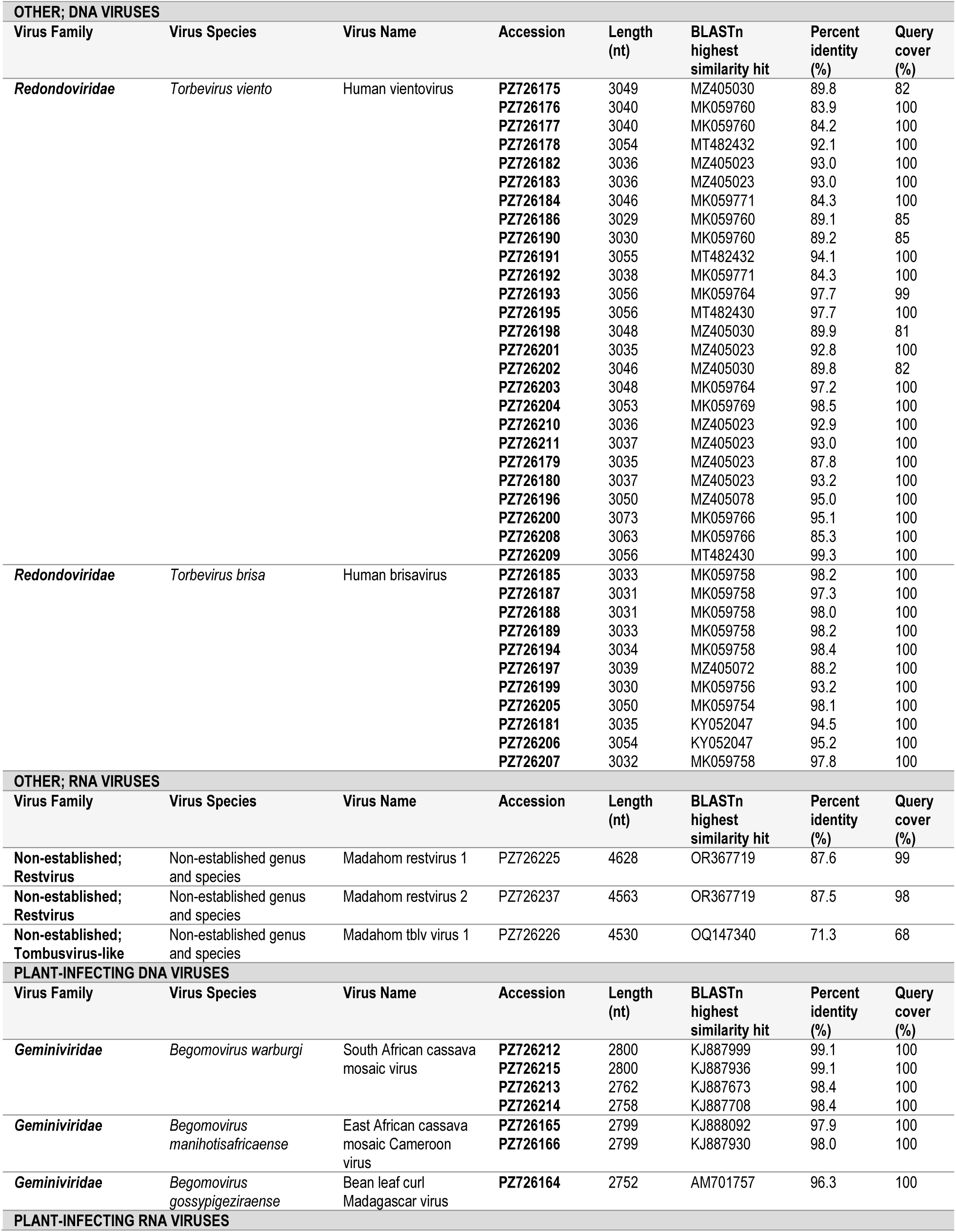
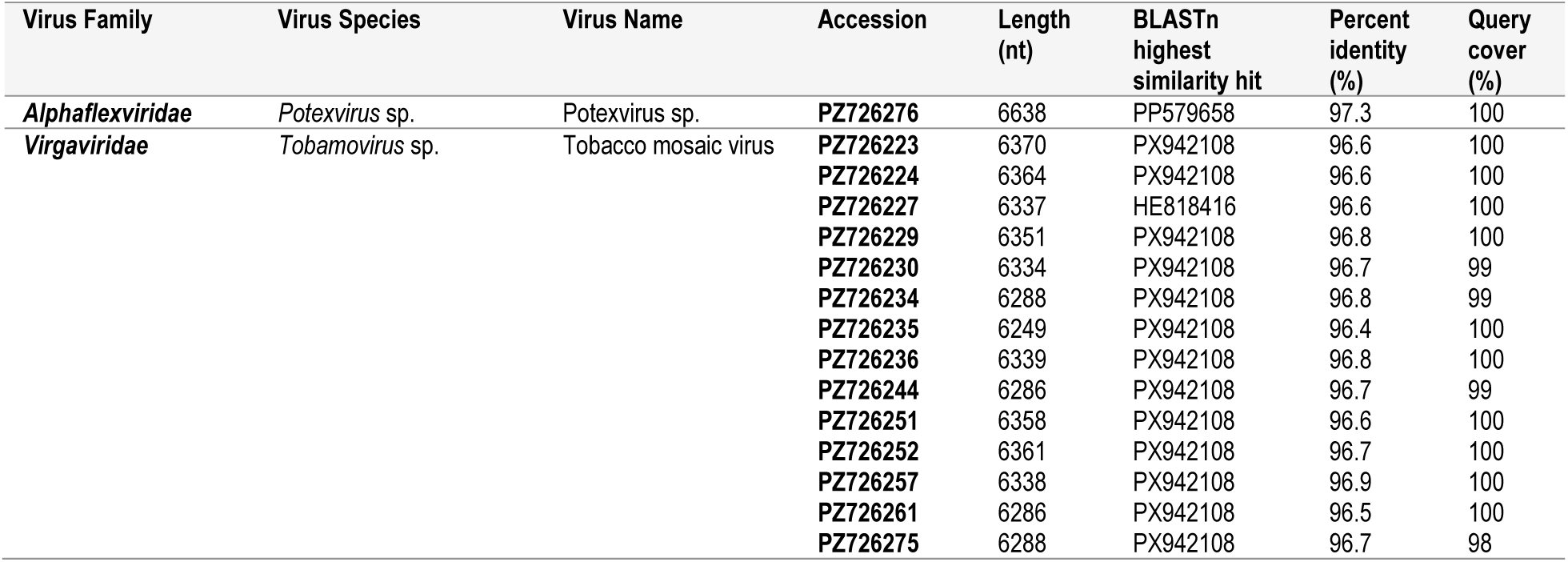
Summary of protist-infecting viruses, plant-infecting viruses, and viruses with unknown hosts identified in oral/nasal swab samples and their similarity to published sequences.

After compiling human-derived viral sequences from Madagascar (NCBI Virus) (Brister et al., 2015), we found that coding-complete sequences were previously available for viruses in the following genera with human-infecting members (Figure 2): *Alphavirus, Betacoronavirus, Bocaparvovirus, Deltavirus, Enterovirus, Lyssavirus, Metapneumovirus, Orthopneumovirus, Orthorubulavirus, Pegivirus, Phlebovirus* (Bird et al., 2007; Carroll et al., 2011; Lockwood et al., 2026; Moyen et al., 2014; Razafindratsimandresy et al., 2014; Sauvage et al., 2021). A vast majority of the available human-derived sequences from Madagascar (Figure 2) were human immunodeficiency virus (HIV) 1 and enterovirus sequences (Rakoto-Andrianarivelo et al., 2008; Rakotomalala et al., 2024; Razafindratsimandresy et al., 2022; Volle et al., 2019). Therefore, this study presents the first publicly accessible coding-complete viral sequences from human communities in Madagascar from multiple DNA viral families, including *Adenoviridae, Anelloviridae, Orthoherpesviridae, Papillomaviridae, Polyomaviridae*, and *Redondoviridae* as well as a few RNA viral families, such as *Astroviridae* and *Sedoreoviridae* (Figure 2). Collectively, these coding-complete viral sequences substantially expand the genomic representation of human-associated viruses from Madagascar and provide a foundation for our understanding of circulating viruses of both public health and surveillance relevance.

**Figure 2.**
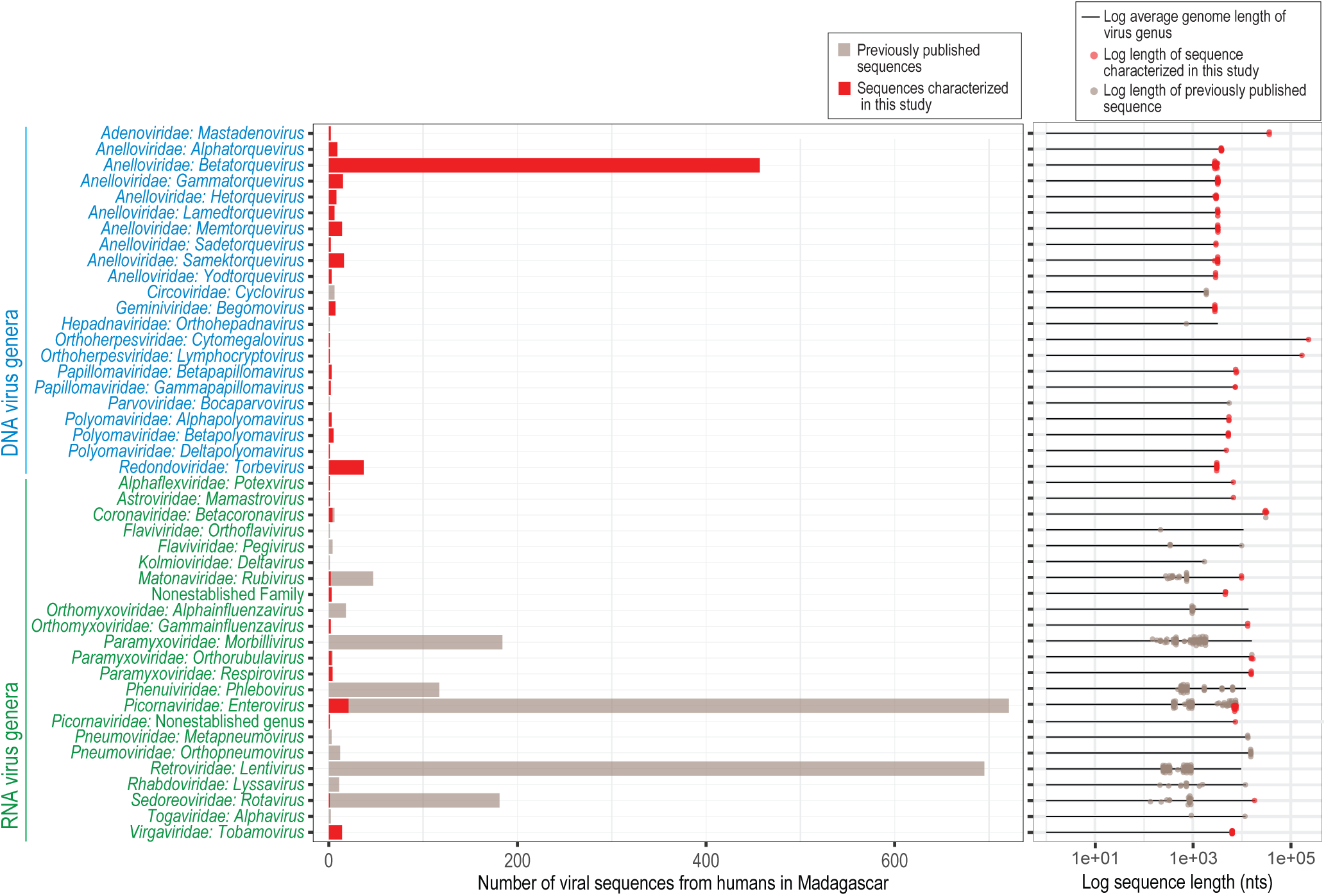
The number of viral sequences previously published (available on NCBI Virus; tan) and viral sequences characterized in this study (red) by genus level (left). The log-scaled lengths of viral sequences previously published (tan) or characterized here (red) were additionally plotted alongside the log-scaled average genome length of the virus genus as represented by a black line, highlighting small partial sequences versus coding-complete sequences for each virus genus (right).

### Pathogenic human-infecting DNA Viruses

Rather than primarily causing acute infection, many human-infecting DNA viruses are known to demonstrate long-term, stable infections spanning months to years to entire lifespans. While many DNA virus infections in healthy humans present asymptomatically, virus pathogenicity and disease severity can be worse in immunocompromised individuals and young children. Complete or coding-complete genomes of viruses in the families *Adenoviridae*, *Orthoherpesviridae*, *Papillomaviridae*, and *Polyomaviridae* were characterized in this study and shared >90% similarity with sequences of viruses commonly circulating in human populations globally (Table 1). These are the first genomes from these families publicly available from people in Madagascar, although some of these viruses have been previously found to have high prevalence in Malagasy communities in urban and rural settings, though no sequence information was available (Bigoni et al., 2018; Garsevanidze et al., 2026; Kutz et al., 2023; Lockwood et al., 2026). While papillomaviruses and polyomaviruses are unlikely to be associated with the febrile symptoms seen among individuals in this study, herpesvirus and adenovirus infections have been associated with fevers and other symptoms (Khanal et al., 2018; Kleinschmidt-DeMasters and Gilden, 2001). The DNA virus sequence data from individuals in rural communities in southeastern Madagascar presented here shows the similarity of human-infecting DNA viromes globally.

The *Papillomaviridae* and *Polyomaviridae* families are composed of oncogenic viruses with circular, dsDNA genomes. In this study, we identified the complete genomes of three known papillomavirus types including human papillomavirus (HPV) 23, 47, and 49 (PZ726167, PZ726168, PZ726169) which are members of the species *Betapapillomavirus 1, Betapapillomavirus 2,* and *Betapapillomavirus 3* (Table 1, Supplementary Table 1). We additionally determined the complete genomes of two non-classified papillomavirus types (PZ726170, PZ726171) that are members of the *Gammapapillomavirus* genus with similarity to HPV-mSK_020 (MH777168) and HPV-mSK_227 (MH777366) which have been identified in healthy human skin samples (USA, 2014-2015) (Tirosh et al., 2018) (Table 1), and thus far have not been associated with a high risk of cancer. For polyomaviruses, we identified multiple complete genomes of polyomaviruses, including Merkel cell polyomavirus (*n* = 3 genomes; PZ726172-PZ726174), WU polyomavirus (*n* = 5; PZ726217-PZ726221), and STL polyomavirus (*n* = 1; PZ726216) (Table 1, Supplementary Table 1). Although WU polyomavirus has been previously reported in Madagascar (Lockwood et al., 2026), this study documents the presence and diversity of Merkel cell (species *Alphapolyomavirus quintihominis*), WU (species *Betapolyomavirus quartihominis*), and STL (species *Deltapolyomavirus undecihominis*) polyomaviruses representing three species across three *Polyomaviridae* genera. Polyomaviruses are associated with a variety of diseases; Merkel cell polyomavirus has been tied to Merkel cell carcinoma, WU polyomavirus has been associated with respiratory symptoms, and STL polyomavirus has an undefined role in disease (Fazeli et al., 2025).

Adenoviruses (family *Adenoviridae*) have been identified not only as pathogens causing respiratory infection but also as pathogens associated with pediatric diarrheal disease (Lee et al., 2020). Two adenovirus genomes were identified here representing genotypes of HAdV-C: human adenovirus 1 (HAdV-C1; PZ726819; species *Mastadenovirus caesari*) and human adenovirus 89 (HAdV-C89; PZ726818; species *Mastadenovirus caesari*). HAdV-C types are primarily associated with respiratory infection and can establish latent infections while persisting in lymphoid tissues. HAdV-C1 is particularly associated with highly prevalent, persistent infections in young children (Akello et al., 2020; Garnett et al., 2002). HAdV-C89 represents a recombinant HAdV-C lineage that thus far has been characterized in samples from China, Kenya, Russia, and the USA, although it is considered a rarer genotype (Dhingra et al., 2019; Hirose et al., 2025; Lambisia et al., 2024; Yolshin et al., 2026).

Viruses in the *Orthoherpesviridae* family have adapted to their hosts over time, demonstrating a variety of mechanisms for immune evasion along with an ability to undergo latency and reactivation (Gatherer et al., 2021). Herpesviruses have large, linear dsDNA genomes ranging in length from ∼120-241 kb usually containing >70 genes (Gatherer et al., 2021). Two herpesvirus genomes were identified in this study representing human cytomegalovirus (HCMV; species *Cytomegalovirus humanbeta5*; PZ726821) within the *Betaherpesvirinae* subfamily and EBV (species *Lymphocryptovirus humangamma4*; PZ726820) within the *Gammaherpesvirinae* subfamily. HCMV was found to be present in seven individuals and EBV in two individuals. For both HCMV and EBV, infection usually presents asymptomatically but illness (e.g., fever, fatigue, blisters, rash, swollen glands, respiratory illness) can be severe in young or immunocompromised individuals, and during pregnancy (Fulkerson et al., 2021).

### Pathogenic human-infecting RNA viruses

In contrast to the human-infecting DNA virome which is comprised of many viruses that commonly establish long-term or latent infections, the human-infecting RNA virome is typically composed of viruses that cause acute infections, and demonstrate high mutation rates, although some RNA virus infections are persistent (Griffin, 2022; Grinde, 2013; Mashhadi Abolghasem Shirazi, 2026). Pathogenic human-infecting RNA viruses impose substantial burdens on public health, particularly in lower-income countries, where they contribute to a range of respiratory, gastrointestinal, neurological, hemorrhagic, developmental and other systemic illnesses. Studies on the human-infecting oral and nasal RNA viromes usually reveal a variety of respiratory viruses (e.g. rhinoviruses, influenza viruses, respiratory syncytial virus, coronaviruses) (Rajagopala et al., 2021; Yao et al., 2025). However, given the extensive diversity of RNA viruses, many studies fail to capture the breadth of viral genetic diversity present in study populations, instead recovering only partial genomic fragments primarily using targeted approaches. Coding-complete human-infecting RNA virus sequences were characterized in this study from seven viral families including *Astroviridae, Coronaviridae, Matonaviridae, Orthomyxoviridae, Paramyxoviridae, Picornaviridae,*and *Sedoreoviridae* (Table 2). Many of these viruses have been previously found to cause an array of clinical manifestations of disease in Malagasy people including fever, rash, respiratory illness, and diarrhea (Hoffmann et al., 2012; Papaventsis et al., 2008; Randremanana et al., 2016; Razafindratsimandresy et al., 2026). The majority of RNA virus sequences characterized in this study shared >90% similarity with known viral sequences, with the exception of one divergent picornavirus (madahom bibyvirus 1) (Table 2).

In the *Matonaviridae* family, we identified two coding-complete rubella virus genomes (PZ726241 and PZ726242; Table 2, Supplementary Table 1). Rubella virus is transmitted through contact with respiratory secretions and infection is characterized by fever and rash, although infection during pregnancy can result in congenital rubella syndrome, stillbirth, or miscarriage (Gregg, 1991; Lambert et al., 2015; Miller et al., 1982). Rubella virus was present in eight samples across six villages spanning age groups <5, 5-10, 10-15, 15-20, and 30-35 years. The rubella virus sequences identified here represent genotype 2B, a genotype previously identified in communities in Madagascar and known to globally circulate (Razafindratsimandresy et al., 2026) (Figure 3). Based on the phylogeny of rubella genomes, the rubella virus sequences characterized here are distinct members of lineage 2B (Figure 3A). An E1 sequence nucleotide phylogenetic tree focused on sequences only from Madagascar revealed a close phylogenetic relationship between the sequences identified in this study and a sequence from Farafangana, the closest city to the villages sampled in this study (∼27 km away) from a gingival swab collected in August 2019 (Razafindratsimandresy et al., 2026) (Figure 3B). Given the availability and global success of safe and highly effective rubella virus vaccines, this finding highlights how metagenomics of circulating human-infecting viruses in at-risk, rural communities can provide guidance on which vaccination programs should be bolstered.

**Figure 3.**
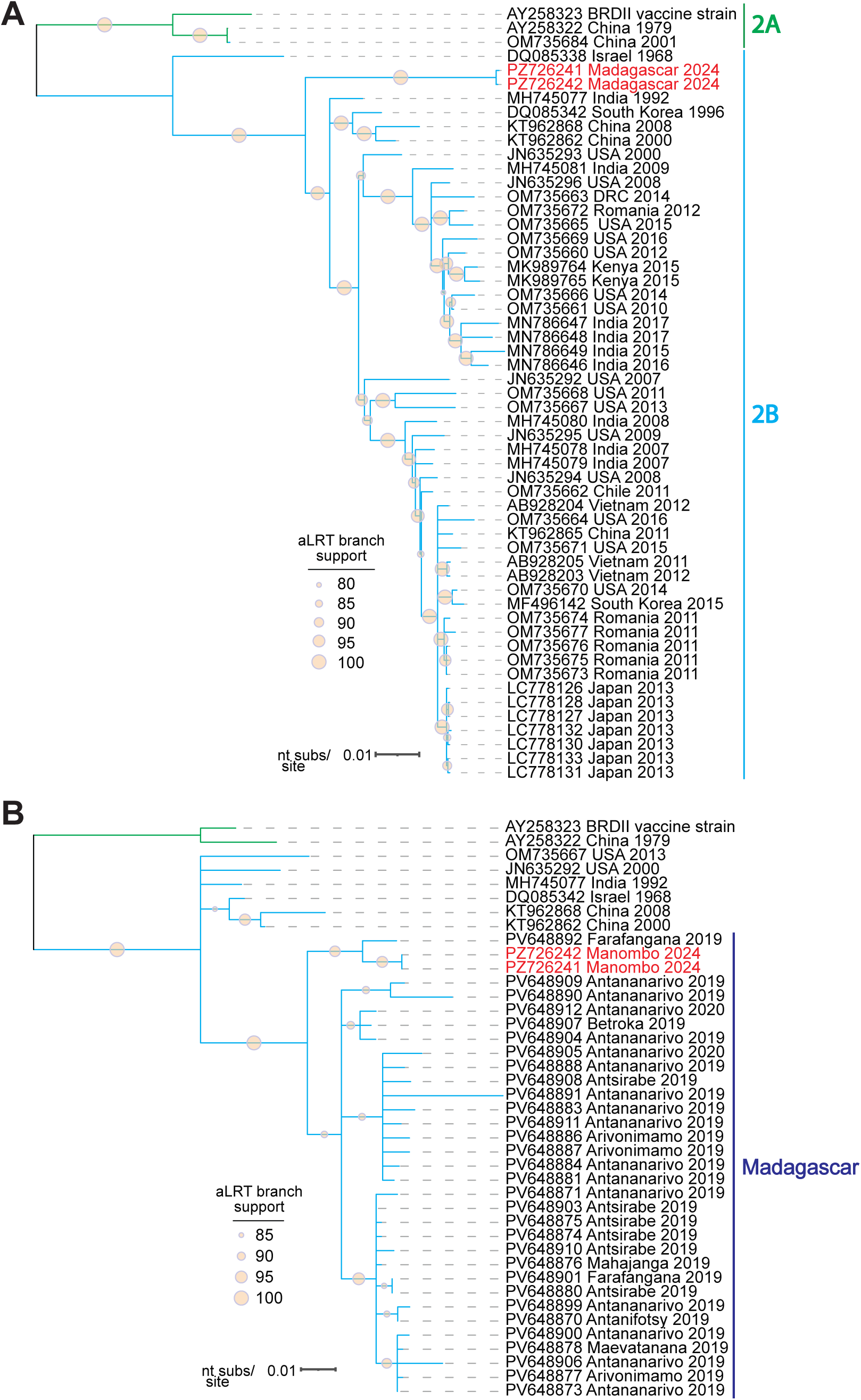
**(A)** Maximum likelihood phylogenetic tree of whole-genome nucleotide sequences for rubella virus in genotype 2B rooted with sequences in genotype 2A. **(B)** Maximum likelihood phylogenetic tree of partial *E1* nucleotide sequences (739 nt) for rubella virus with a focus on available sequences from Madagascar. Viruses characterized in this study are shown in red font.

Influenza C virus (family *Orthomyxoviridae*, genus *Gammainfluenzavirus*) was identified in two individuals aged <5 years from the same village. The influenza C virus genome is composed of seven ssRNA(-) segments totaling ∼10 kb which we were able to recover for both individuals (PZ716971-PZ716984) (Table 2). While the impacts of influenza A and B viruses on Malagasy populations have been more thoroughly studied (e.g. high mortality associated with A(H1N1) in Antananarivo in 2009 (Rajatonirina et al., 2013a), reemergence of A(H1N1), A(H3N2), and B/Victoria following the onset of the SARS-CoV-2 pandemic (Razanajatovo et al., 2025)), studies on the circulation of influenza C virus in Madagascar have been neglected, likely because it causes less severe disease. Given research in Madagascar on the introduction of influenza A(H1N1)pdm09 in 2009 through sporadic initial cases originating from India, Mauritius, and La Réunion (Rajatonirina et al., 2012), we performed BEAST analysis to understand the potential origins of the influenza C virus strain sequenced in this study. Phylogenetically, the Madagascar sequences are a distinct lineage nested within a broader mixed-location clade composed of sequences from Australia, Austria, Cameroon, Hong Kong, and Japan (Figure 4A). We estimate that this influenza C virus strain was introduced to Madagascar around 2009 from Asia, placing its introduction on a similar timeline to influenza A(H1N1)pdm09 (Figure 4A). The Skygrid reconstruction suggests a period of demographic expansion of influenza C viruses during the 1990s, followed by a broadly stable but fluctuating effective population size and a recent decline characterized by substantial uncertainty (Figure 4B).

**Figure 4.**
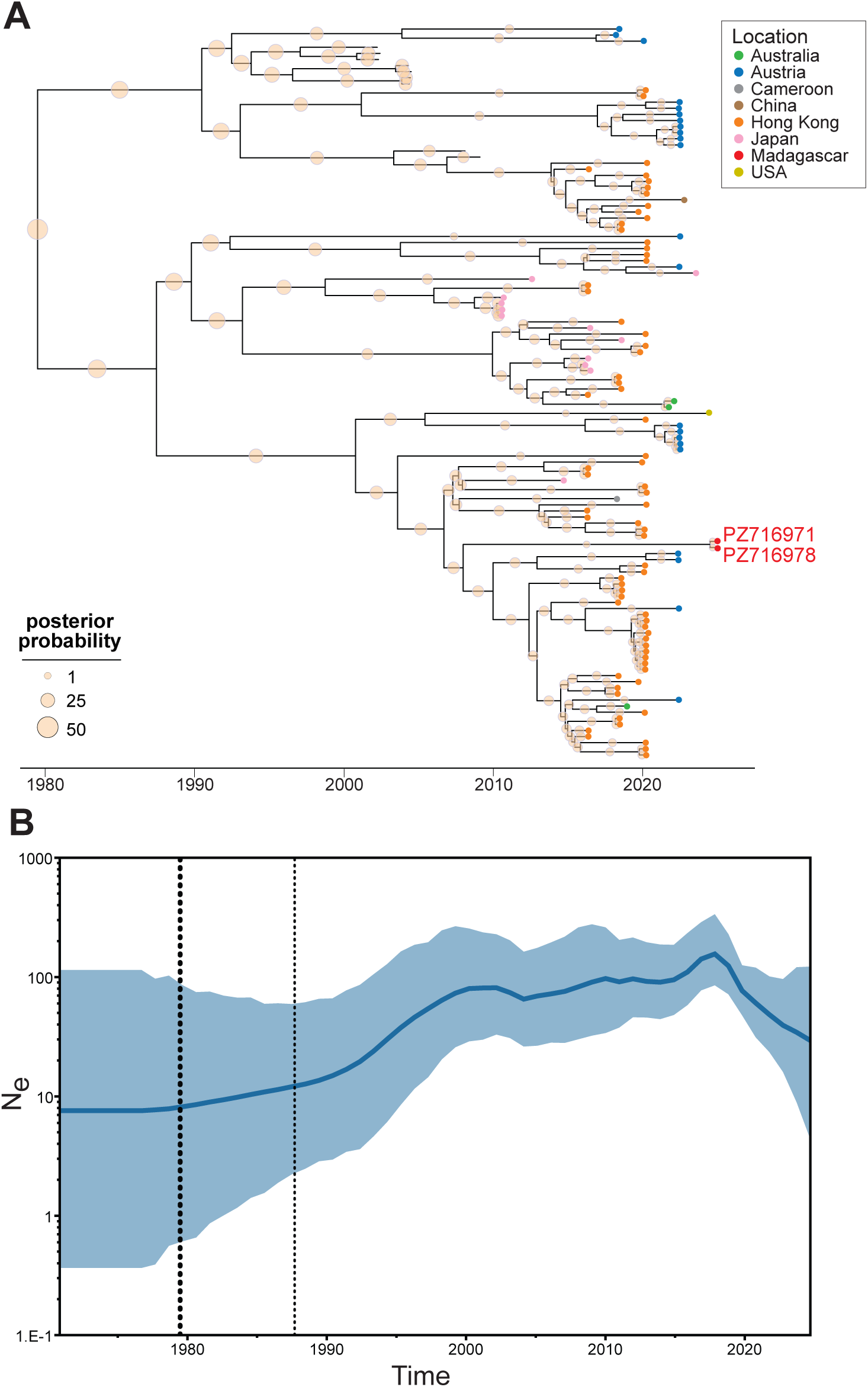
**(A)** Time-scaled Bayesian maximum clade credibility tree of 120 influenza C genomes. **(B)** Bayesian Skygrid reconstruction of the effective population size of influenza C viruses over time.

Within the *Enterovirus* genus in the *Picornaviridae* family, we identified 21 coding-complete virus sequences belonging to five established species: *Enterovirus alphacoxsackie* (n = 1 genome; PZ726267), *Enterovirus betacoxsackie* (n = 1; PZ726272), *Enterovirus coxsackiepol* (n = 3; PZ726222, PZ726250, PZ726253), *Enterovirus alpharhino* (n = 10; PZ726238, PZ726239, PZ726246, PZ726248, PZ726254, PZ726256, PZ726262, PZ726263, PZ726268, PZ726270), and *Enterovirus cerhino* (n = 6) (PZ726231, PZ726255, PZ726258–PZ726260, PZ726273) (Table 2, Supplementary Table 1). Many of these enteroviruses are genetically similar to common, globally circulating types, including Coxsackievirus A6 and B4 and rhinovirus A1, A19, A20, A45, A77, A89, C2, C7, and C10. Rhinovirus C2 and A20 had the highest detection frequencies (9% and 5%, respectively) in children aged <5 years and Coxsackievirus A6 and Coxsackievirus B4 were each detected in a single child in the <5 year age group. While rhinoviruses generally cause mild common cold symptoms, Coxsackievirus A6 infection is considered a primary cause of hand, foot, and mouth disease (HFMD) (Bian et al., 2015), and Coxsackievirus B4 has been associated with a variety of respiratory, pancreas, heart, and central nervous system diseases, although it has also been associated with HFMD (Xiao et al., 2021). These enteroviruses share 94.5-99.3%coding complete genome nucleotide pairwise identity with known rhinovirus and coxsackievirus sequences (Table 2).

In addition to coxsackieviruses and rhinoviruses, we identified two enterovirus genotypes C105 and C117 which are considered rare but emerging genotypes. While first characterized in a patient with acute flaccid paralysis in 2010 in the Democratic Republic of the Congo (Lukashev et al., 2012), enterovirus C105 was also identified in samples collected in 2010 in Burundi, Italy, Peru, and Romania and was subsequently detected in various European, Asian, Oceanic, and North American countries from 2011-2025 (Fall et al., 2025; Fernandez-Garcia et al., 2025; Horner et al., 2015; Li et al., 2019; Majumdar et al., 2021; Piralla et al., 2015; Richter et al., 2013; Todd et al., 2015; Tokarz et al., 2013). Enterovirus C105 has been tied to both neurological and respiratory symptoms (Fall et al., 2025; Horner et al., 2015; Lukashev et al., 2012). Phylogenetic analysis of enterovirus C105 VP1 nucleotide sequences revealed that the sequences detected here in Malagasy individuals are most closely related to sequences identified in European (France, Italy, Spain) and North American (USA) samples collected in 2023-2024, although this lineage additionally includes a sequence from a sample collected in Spain in 2019 (Figure 5A). Enterovirus C117, an emerging type putatively associated with respiratory illnesses, was initially characterized in samples from Lithuania (2011), China (2007-2017), and Mongolia (2011-2013), but was subsequently identified in the USA, Netherlands, Cameroon, and France (Daleno et al., 2012; Du et al., 2019; Kenmoe et al., 2021; Van Leer-Buter et al., 2016; Xiang et al., 2014). Using VP1 sequences, the enterovirus C117 sequence identified here shares the closest phylogenetic relationship with a sequence in a sample collected in China (2017; MT338567) (Figure 5B). The diversity of picornavirus sequences characterized here suggests the need for increased monitoring of enterovirus types and associated symptoms, from HFMD to acute flaccid paralysis, in Malagasy populations.

**Figure 5.**
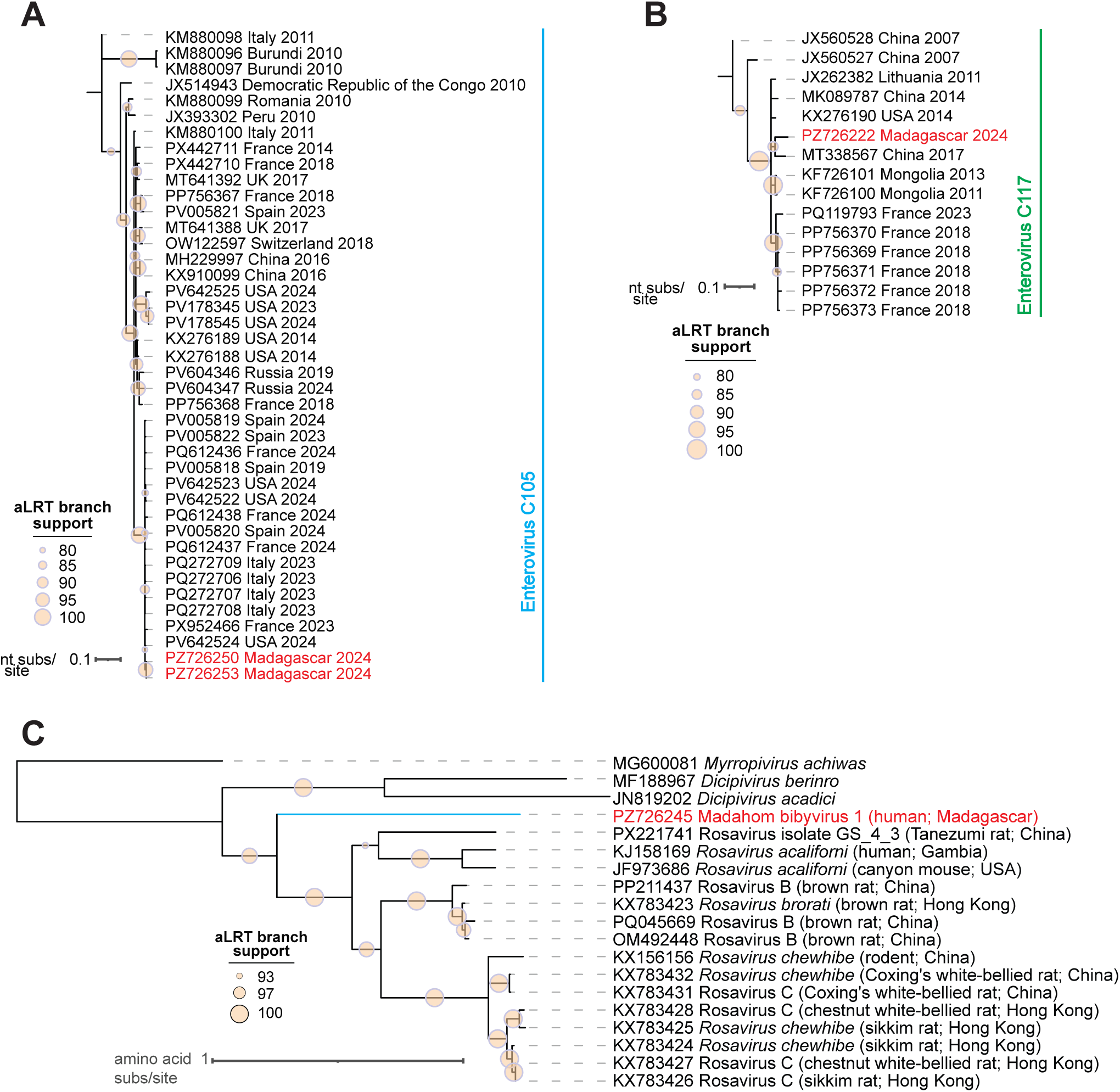
**(A)** Maximum likelihood phylogenetic tree of VP1 nucleotide sequences of Enterovirus C105, **(B)** Maximum likelihood phylogenetic tree of VP1 nucleotide sequences of Enterovirus C117. **(C)** Maximum likelihood phylogenetic tree of polyprotein amino acid sequences of picornaviruses (genera *Rosavirus, Dicipivirus, Myrropivirus*) related to madahom bibyvirus 1. Viruses characterized in this study are shown in red font.

Further, in the sample of one child aged <5 years, we identified a highly divergent picornavirus sequence which we have named madahom bibyvirus 1 (PZ726245), that shows a distant relationship to rosaviruses, though it likely represents a putative new genus (Table 1, Figure 5C). Rosaviruses have primarily been identified in samples from multiple rodent species and shrews, with the exception of rosavirus A2 (species *Rosavirus acaliforni*) which was detected in fecal samples from children in Gambia (Lim et al., 2014; Zell et al., 2017; Zhang et al., 2024). Although rosavirus transmission and pathogenicity remain understudied, rosavirus C has been found to cause multisystemic diseases in a mouse model (Lau et al., 2016) and rosavirus genotype B has been detected in rat spleens, demonstrating pathogenic potential in mammals (Boros et al., 2019). Because it was only detected in one human oral/nasal swab, we cannot confirm whether madahom bibyvirus 1 is actively infecting humans versus environmentally-derived, but its phylogenetic position near putatively mammal-infecting genera *Rosavirus* and *Dicipivirus* demonstrates mammal-infecting potential.

Within the *Betacoronavirus* genus of the *Coronaviridae* family, three coding-complete human coronavirus OC43 sequences (species *Betacoronavirus gravedinis*; PZ726240, PZ726249, PZ726265) and one human coronavirus HKU1 sequence (species *Betacoronavirus hongkongense*; PZ726233) were characterized (Table 2, Supplementary Table 1). Human coronavirus OC43 and HKU1 primarily replicate in upper respiratory tract epithelial cells and usually cause mild respiratory infections (Liu et al., 2021). Human coronavirus OC43 was present in eight individuals from six villages in age groups <5, 5-10, 15-20, and 20-25 years. Human coronavirus HKU1 was present in six individuals from five villages in age groups <5, 5-10, 20-25, 30-35, and 40-45 years. Human coronavirus OC43 sequences shared the highest genome-wide pairwise identity with genomes from the USA (99.9%) that were sampled in March 2024 (PV685491, PV685491) (Table 2). The human coronavirus HKU1 sequence shared the highest genome-wide pairwise identity (99.9%) to a genome from the USA collected in February 2024 (PV685467; Table 2). Additionally, although we were only able to assemble fragments (78.2% genome coverage, 1719 reads), SARS-CoV-2 was present in one individual aged 5-10 years. A complete nucleocapsid phosphoprotein was obtained and demonstrated 100% nucleotide identity to sequences from the USA sampled in June 2024 (PQ040855, PP819325). These results highlight the circulation of betacoronaviruses in southeastern Madagascar and bring attention to the close genetic similarity of the coronaviruses detected in this study from samples collected in September 2024 to globally circulating variants identified from February-June 2024 indicating a recent introduction.

In the *Paramyxoviridae* family, we identified three coding-complete sequences of human parainfluenza virus type 2 (HPIV-2; species *Orthorubulavirus laryngotracheitidis*; PZ726228, PZ726243, PZ726274) and five coding-complete sequences of human parainfluenza virus type 3 (HPIV-3; species *Respirovirus pneumoniae*; human respirovirus 3; PZ726232, PZ726247, PZ726264, PZ726266, PZ726269) (Table 2, Supplementary Table 1). HPIVs replicate in lower and upper respiratory tract epithelial cells and primarily cause mild respiratory infections similar to the common cold, although laryngotracheitis, bronchiolitis, and pneumonia are possible (Kennedy et al., 2017; Pawełczyk and Kowalski, 2017; Zhu et al., 2024). HPIV-2 and −3 have been found to be prevalent in children experiencing febrile illness or respiratory infection in rural communities in Madagascar (Hoffmann et al., 2012; Maeder et al., 2016). Here, HPIV-2 was present in five samples across four villages spanning age groups <5, 5-10, 15-20, and 20-25 years. The HPIV-2 sequences identified here are within clade G1, with two sequences representing clade G1A (PZ726228, PZ726274) and one sequence representing clade G1C (PZ726243), showing the co-circulation of at least two HPIV-2 lineages in these communities (Supplementary Figure 2A). HPIV-3 was present in six samples across three villages spanning age groups <5 and 20-25 years. The HPIV-3 sequences characterized here (PZ726269, PZ726247, PZ726264, PZ726232, PZ726266) are positioned in clade C3A, one of the primary HPIV-3 lineages that is circulating globally (Mansour et al., 2025; Zhu et al., 2024) (Supplementary Figure 2B).

Rotaviruses (family *Sedoreoviridae*) and astroviruses (family *Astroviridae*) are enteropathogens linked to pediatric gastroenteritis and severe diarrheal disease and dehydration, including in Malagasy children (Papaventsis et al., 2008; Randremanana et al., 2016; Razanajatovo et al., 2023). Rotaviruses possess segmented genomes composed of 11 segments of dsRNA totaling ∼18.5 kb in length, whereas astroviruses have non-segmented ssRNA(+) genomes ∼6.8 kb in length. A rotavirus genome (11 segments; PZ726277 - PZ726287) was identified in an individual aged 20-25 years (Table 2, Supplementary Table 1). The rotavirus genome represents rotavirus A and genotype constellation G12-P[6]-I1-R1-C1-M1-A1-N1-T1-E1-H1, showing a Wa-like backbone. Based on the maximum likelihood phylogenetic tree of VP4 nucleotide sequences focused on the P[6] genotype, the rotavirus VP4 sequence characterized here is positioned closely with rotavirus sequences in stool samples from Pakistan from 2019-2023 (PV768348, PV768352, PV768353 (Usman et al., 2024)) and is not positioned with P[6] sequences from Madagascar from 2008-2009. Previous studies in Madagascar from 2004-2005 and 2008-2009 characterized G1, G3, G4, and G9 genotypes for VP7 and P[6], P[8], and P[9] genotypes for VP4 with G9 and P[8] most commonly detected (Adiku et al., 2010; Razafindratsimandresy et al., 2013). However, additional studies on rotavirus A in Madagascar from 2013-2016 detected G3-P[8] and G2-P[4] as the most common genotypes, documenting a shift after rotavirus vaccine introduction in 2014 (Rahajamanana et al., 2018; Seheri et al., 2018). Although reports in Madagascar remain limited, G12-P[6] has been increasingly documented in various African countries including Ethiopia, South Africa, Togo, Uganda, Zambia, and Zimbabwe, though G12 has been globally circulating both pre- and post-vaccine introduction (Mokoena et al., 2021; Rakau et al., 2021). Thus, our findings geographically expand the surveillance of G12-P[6] in African countries to Madagascar. A human astrovirus genome (PZ726271) was characterized in a child in the <5 year group (Table 2, Supplementary Table 1). The astrovirus genome identified here belongs to human astrovirus type 3 (HAstV-3) and is phylogenetically positioned in a lineage containing HAstV-3 sequences from Kenya (MW485043), Malawi (PX255447), South Africa (MT267482), and China (OM468668, OM468669) (2015-2019), suggesting a history of HAstV-3 transmission between African countries.

### Pathogen co-occurrences

While many of the viruses identified here (e.g. polyomaviruses, herpesviruses, rhinoviruses) primarily present asymptomatically or cause mild symptoms in healthy hosts, we emphasize that co-infections and factors associated with socioeconomic and environmental conditions in the region (e.g., malnutrition, anemia, vector exposure and *Plasmodium* infection) can weaken host immune systems and worsen symptoms associated with viral or protozoan infection, potentially resulting in more severe disease (e.g., Merkel cell carcinoma, Burkitt lymphoma, pneumonia). Pathogen co-infections have also been found to result in a higher chance of mortality, notably among young children in Madagascar (Rajatonirina et al., 2013b). We detected at least one pathogenic DNA virus in 22 individuals, and at least one pathogenic RNA virus in 61 individuals Among these, pathogenic DNA virus-DNA virus, DNA virus-RNA virus, and RNA virus-RNA virus co-occurrences were seen in two, six, and five individuals, respectively (Figure 7).

**Figure 6.**
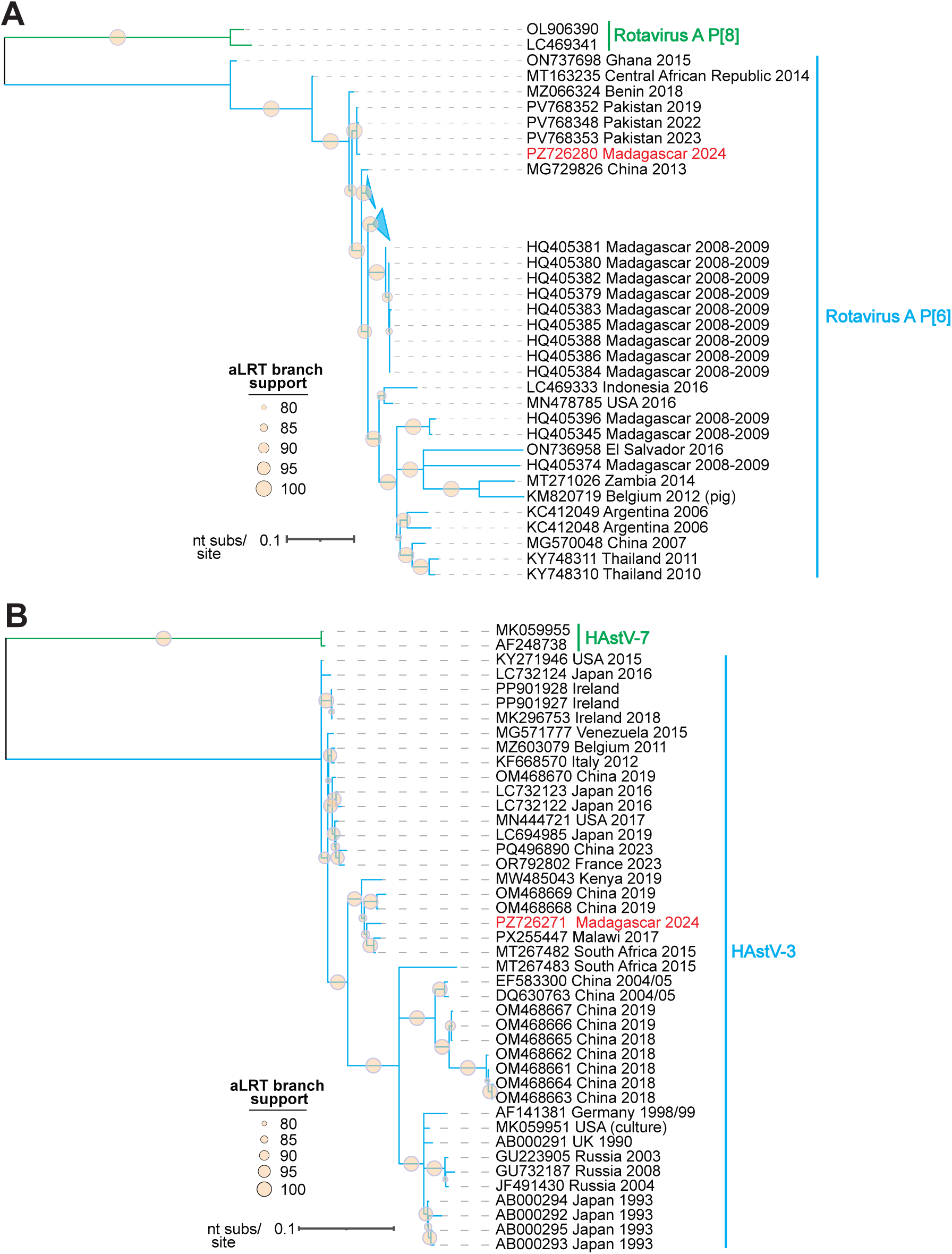
**(A)** Maximum likelihood phylogenetic tree of *VP4* nucleotide sequences of rotavirus A genotype P[6] with a focus on sequences from Madagascar rooted with sequences of rotavirus A genotype P[8]. **(B)** Maximum likelihood phylogenetic tree of *orf2* nucleotide sequences of human astrovirus type 3 sequences rooted with sequences of human astrovirus type 7. Viruses characterized in this study are shown in red font.

**Figure 7.**
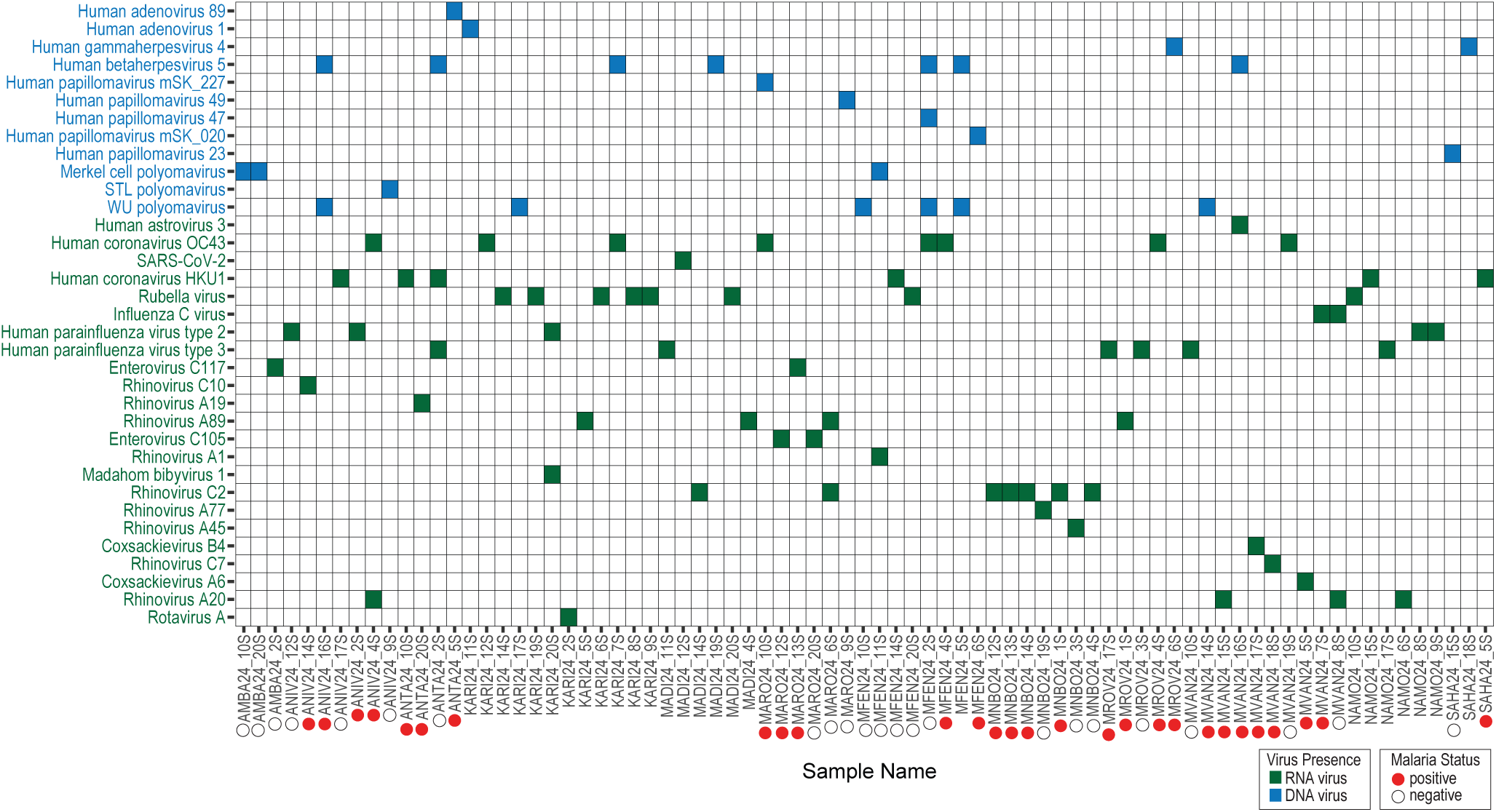
Co-occurrence of pathogenic DNA and RNA viruses across samples. Malaria status was available for a subset of samples. Note that madahom bibyvirus 1 is not confirmed to be a pathogenic human-infecting virus.

Virus co-occurrences within the respiratory tract may impact disease severity. Here, we document influenza C virus-rhinovirus A20 (MVAN24_8S), human coronavirus OC43-rhinovirus A20 (ANIV24_4S), and rhinovirus A89-rhinovirus C2 (MARO24_6S) co-occurrences all involving RNA viruses with overlapping respiratory tract epithelial cell tropism (Figure 7). We also see co-occurrences of viruses known to cause lasting, latent infections, such as human cytomegalovirus (HCMV)-WU polyomavirus co-occurrence in three individuals, which has been documented previously in various populations (Babakir-Mina et al., 2009; Mueller et al., 2009). HCMV presence was usually accompanied by other pathogens including *P. falciparum*, human astrovirus, coronaviruses, papillomaviruses, polyomaviruses, and paramyxoviruses with one individual infected with viruses across four families: HCMV (*Orthoherpesviridae*), HPV type 47 (*Papillomaviridae*), WU polyomavirus (*Polyomaviridae*), and human coronavirus OC43 (*Coronaviridae*) (Figure 7).

Aided by our RDT testing, we also documented virus-protozoan co-occurrences (Figure 7). For individuals where malaria RDT status was available, viruses in DNA families *Orthoherpesviridae, Papillomaviridae,* and *Polyomaviridae* and RNA families *Paramyxoviridae*, *Picornaviridae*, and *Orthomyxoviridae* were identified in the samples of both *P. falciparum*-positive and *P. falciparum*-negative individuals (Figure 7). We identified EBV in a *P. falciparum*-positive individual, a notable virus-protozoan co-occurrence given that *P. falciparum* impairs immune control of EBV infection while promoting the expansion of EBV-infected B cells, which increases the risk of Burkitt lymphoma (Moormann and Bailey, 2016). Yet, Burkitt lymphoma cases are rare, and the mobile healthcare clinic has not reported any cases of Burkitt lymphoma, despite high prevalence of *P. falciparum* in these communities.

Because malaria suppresses the immune system, we expected to see a relationship between malaria status and pathogenic virus detection. However, using a generalized linear mixed model controlling for village as a random effect, malaria status was not significantly associated with pathogenic virus detection (estimate = −0.030, se = 0.364, p = 0.934). The absence of an observed relationship may reflect the focus of our study on samples from the oral and nasal cavities. We found that males trended towards higher odds of pathogenic virus detection (estimate = 0.653, se = 0.368, p = 0.076), although these associations were not significant. However, age was significantly associated with pathogenic virus detection (estimate = −0.024, se = 0.010, p = 0.017). This suggests that younger individuals had higher odds of pathogenic virus detection likely reflecting acquired natural immunity in older individuals.

The contributions of co-occurring viral and protozoan pathogens in individuals in this study to the febrile illness symptoms recorded cannot be confidently parsed. However, in *P. falciparum*-negative individuals, we suspect that viral co-infections or infections from pathogens including rubella virus, coronaviruses, and picornaviruses were likely causing symptomatic disease.

### Other human-associated viruses

While known pathogenic human-infecting viruses are often studied for their direct roles in disease, the human virome consists of various other viruses including non-pathogenic human-infecting viruses providing insight into host immune systems (e.g. anelloviruses and pegiviruses), microbiome-associated viruses (e.g. redondoviruses and bacteriophages), or diet-associated viruses (e.g. geminiviruses and tobamoviruses). Metagenomic analysis of oral and nasal swabs provide an opportunity to detect these additional viruses.

#### Anelloviruses

Anelloviruses are single-stranded, animal-infecting DNA viruses exhibiting extensive genetic diversity, high prevalence, and frequent co-infections in a host (Kaczorowska and Van Der Hoek, 2020; Varsani et al., 2021). Although considered non-pathogenic and potentially commensal, anelloviruses have been found to interact with the human immune system, as increased anellovirus replication and load has been reported for some initial studies on immunocompromised groups and higher anellovirus prevalence has been documented in febrile individuals (De Vlaminck et al., 2013; McElvania TeKippe et al., 2012; Thom and Petrik, 2007). However, the host and environmental factors shaping anellovirus diversity and dynamics remain poorly understood, particularly for populations experiencing significant immunological pressures, such as malaria or other pathogens, as well as conditions associated with high levels of poverty and malnutrition. Anellovirus taxonomic composition, primarily in blood samples, has been found to differ across age groups with some studies finding betatorqueviruses and gammatorqueviruses dominating in infants followed by a shift to alphatorqueviruses dominating in most adults (Kaczorowska et al., 2022; Li et al., 2022; Timmerman et al., 2025). A study on blood samples from febrile Tanzanian children aged two months to five years found the highest detection frequency of betatorqueviruses, although it is unclear if this was maintained across ages (Cordey et al., 2021). As most studies on anellovirus dynamics have focused on blood samples, trends in anellovirus genus composition, richness, and co-occurrence in the upper respiratory tract remain largely uncharacterized.

Here, using oral and nasal swabs, we identified complete anellovirus genomes (PZ726288–PZ726817) that are members of all nine established human-infecting anellovirus genera: *Alphatorquevirus* (*n* = 9 genomes, 8 vOTUs; detection frequency 27%), *Betatorquevirus* (*n* = 457 genomes, 395 vOTUs; detection frequency 50%), *Gammatorquevirus* (*n* = 15 genomes,11 vOTUs; detection frequency 17%), *Hetorquevirus* (*n* = 8 genomes, 5 vOTUs; detection frequency 5%), *Lamedtorquevirus* (*n* = 6 genomes, 4 vOTUs; detection frequency 5%), *Memtorquevirus* (*n* = 14 genomes, 11 vOTUs; detection frequency 10%), *Sadetorquevirus* (*n* = 2 genomes, 1 vOTU; detection frequency 1%), *Samektorquevirus* (*n* = 16 genomes, 9 vOTUs; detection frequency 15%), and *Yodtorquevirus* (*n* = 3 genomes, 2 vOTUs; detection frequency 2%) (Supplementary Table 2). Twenty-four samples had vOTUs from *Alphatorquevirus*, *Betatorquevirus*, and *Gammatorquevirus*. Four samples had vOTUs from six different genera.

Anellovirus genus composition was relatively consistent across age, malaria status, and sex with a few notable exceptions (Figure 8A). In infants aged seven months to one year, anellovirus communities were less diverse than other age groups, composed of only betatorqueviruses, gammatorqueviruses, hetorqueviruses, and samektorqueviruses, demonstrating a lack of alphatorquevirus detection, although sample size was low (*n* = 5). Further, average anellovirus vOTU detections and anellovirus diversity were low in individuals aged 30-40 in both males (*n* = 6) and females (*n* = 16). Males had the highest average anellovirus vOTU detections per sample for age groups 15-20 (*n* = 6) and 20-30 (*n* = 8). In stark contrast with the dominance of alphatorqueviruses in some studies on human blood samples, betatorqueviruses dominated across all age groups in this dataset (Figure 8A).

**Figure 8.**
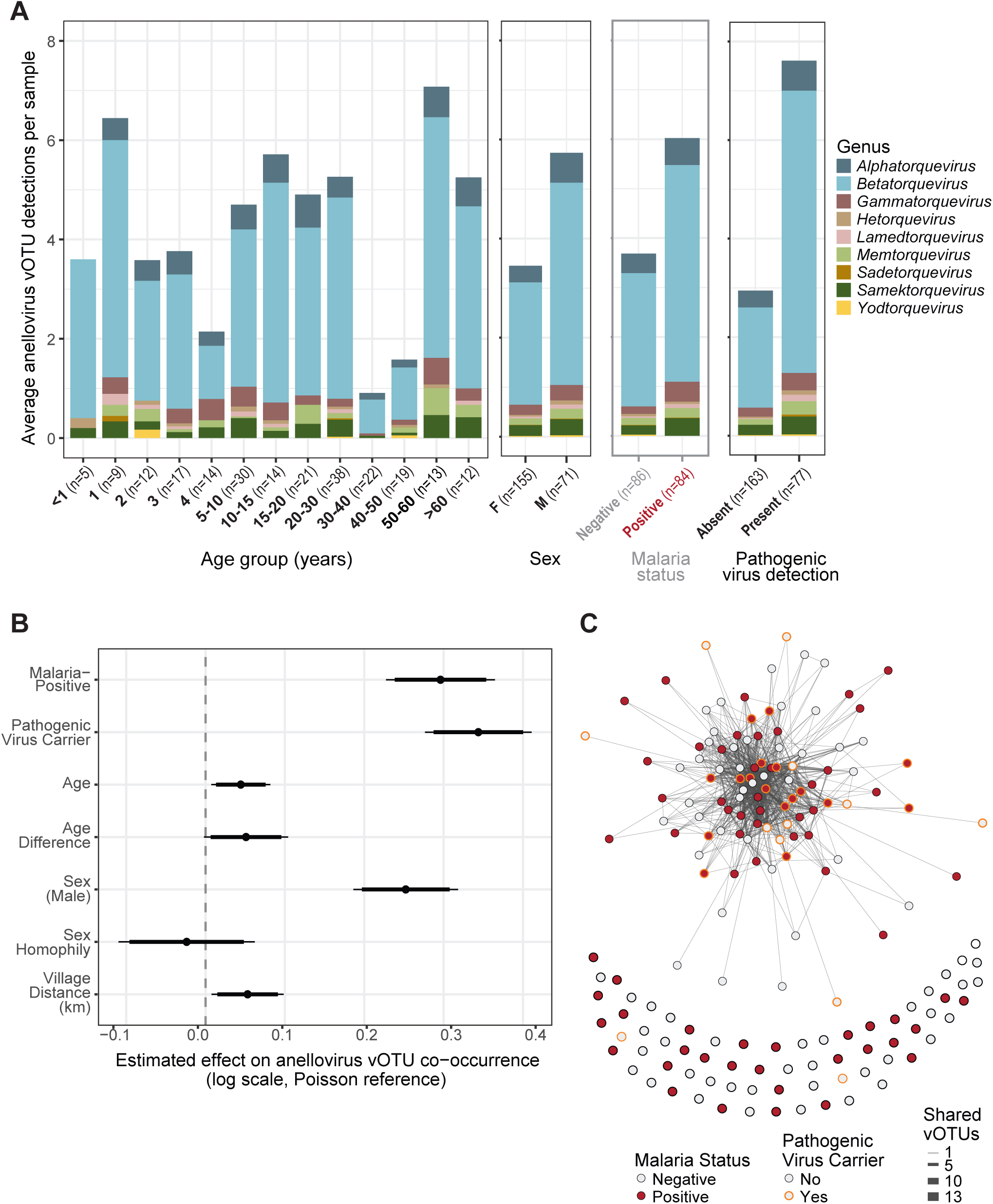
**(A)** Stacked barplot of average anellovirus vOTU detections per sample across age groups, sex, malaria status, and pathogenic virus presence, separated by anellovirus genus. **(B)** ERGM coefficient plot showing host traits associated with virus sharing. Note that the plot excludes the two network structural terms included in the model: “sum” and “nonzero”. Effect sizes are on the log-linear scale (i.e., for every 1 unit increase in a given variable, the number of shared viruses multiplies by e^b, where b is the estimated effect size; because the model includes a “nonzero” term, this rate does not directly represent the number of shared viruses). Positive coefficients represent that a given variable is associated with more shared viruses. **(C)** Co-occurrence network representing humans (nodes) and the anellovirus vOTUs they share with other individuals (edges). Edges are weighted by the number of shared viruses. Isolates (individuals without co-occurring anelloviruses with any other person in the sample; n = 68) are shown.

Intraindividual anellovirus vOTU richness ranged from 0 to 70. Using a generalized linear mixed model controlling for village as a random effect, anellovirus richness was significantly higher in males (estimate = 0.759, se = 0.087, p = <2e-16), younger individuals (estimate = −0.006, se = 0.002, p = 0.007), and those with a positive malaria status (estimate = 0.206, se = 0.090, p = 0.022) or a detected pathogenic virus (DNA or RNA) (estimate = 0.989, se = 0.081, p = <2e-16). Additionally, using an ERGM, we tested the predictive effects of age, age difference, sex, village distance, pathogenic virus detection (DNA or RNA), and malaria status covariates on anellovirus vOTU co-occurrence (n = 170 individuals) (Figure 8B, C). We estimated that people shared 4.8% more anellovirus vOTUs per additional year of age (estimate = 0.05, p = 0.013). Because this finding disagrees with the results of the regression model, we additionally looked at anellovirus vOTU co-occurrence using the broader dataset of 226 individuals for which demographic data was available, excluding malaria (due to gaps in malaria status data) and pathogenic virus status. Similarly to the regression model, we found that younger individuals shared more vOTUs, suggesting a relationship between malaria status and age; we see a malaria prevalence of 68.3% in individuals under 10 years of age and 39.1% in individuals over 10 years of age (Supplementary Figure 3). We recognize that this difference may be the result of higher parasitemia in children resulting in higher detectability via the RDT and missed RDT detection in adults with lower parasitemia due to previous exposure. Pairs farther apart in age tended to share more anelloviruses than individuals close in age (estimate = 0.050, p = 0.069; marginally significant) (Figure 8B), which may reflect cumulative exposure to anelloviruses over time, as well as potential household transmission. Males were estimated to have 28.4% more shared anellovirus vOTUs than females (estimate = 0.250, p <0.001), likely due to mobility across the landscape, but we did not detect evidence for a sex homophily (estimate = −0.023, p = 0.593) (Figure 8B). Individuals with malaria tended to have 34.8% more shared anellovirus vOTUs with others (estimate = 0.299, p < 0.001) compared to individuals testing negative for malaria. Similarly, those with a pathogenic virus tended to share 41.8% more anellovirus vOTUs (estimate = 0.350, p < 0.001) than individuals without pathogenic viruses. The effects of pathogens, both protozoa and viruses, on anellovirus vOTU sharing demonstrates that if the host immune system is impacted by such infections it is likely reflected in anellovirus sharing and persistence resulting in higher richness among individuals in these communities.

### Diet and microbiome related viruses: Redondoviruses, tombus-like viruses, and plant-infecting viruses

Viruses in the *Redondoviridae* family are predicted to infect *Entamoeba* associated with periodontitis in humans (Abbas et al., 2021; Abbas et al., 2019; Keeler et al., 2023; Kinsella et al., 2022). These viruses thus signify oral protist infection. We identified 26 genomes of redondoviruses in the species *Torbevirus viento* (PZ726181, PZ726185, PZ726187–PZ726189, PZ726194, PZ726197, PZ726199, and PZ726205–PZ726207) and 11 genomes of redondoviruses in the species *Torbevirus brisa* (PZ726175–PZ726180, PZ726182–PZ726184, PZ726186, PZ726190–PZ726193, PZ726195–PZ726196, PZ726198, PZ726200–PZ726204, and PZ726208–PZ726211) (Table 3, Supplementary Table 1). Vientoviruses and brisaviruses identified in this study are dispersed across the redondovirus sequence phylogenetic tree and nested among sequences from Africa, Asia, Europe, North America, and South America (Abbas et al., 2019; Cui et al., 2017; Kraberger et al., 2019; Taylor et al., 2021; Thieulent et al., 2025; Tochetto et al., 2021), with the exception of one divergent lineage in *Brisavirus* genus (PZ726197) and one divergent lineage in *Vientovirus* genus (PZ726175, PZ726198, PZ726202) (Supplementary Figure 4).

We identified three virus sequences in tombus-like virus clades which have yet to be classified and assigned a family (Table 3, Supplementary Table 1). Two of these sequences (PZ726237, PZ726225) show similarity to two virus groups that have been primarily identified from metagenomic or metatranscriptomic datasets from humans and other animals, stool-associated tombus-like viruses termed statoviruses (Hause et al., 2021; Janowski et al., 2017; Ramesh et al., 2021; Schneider-Crease et al., 2022) and respiratory-associated tombus-like viruses termed restviruses or statovirus-like viruses (Song et al., 2024). Statoviruses have been identified in the feces of primates (humans, geladas, rhesus macaques, southern pig-tailed macaques), ungulates (cattle, pigs), and rodents (mice) and restviruses have been characterized from human oropharyngeal swabs, bronchoalveolar lavage fluid, and sputum (Hause et al., 2021; Janowski et al., 2017; Ramesh et al., 2021; Schneider-Crease et al., 2022; Song et al., 2024).

The sequences identified here in oral/nasal swabs (PZ726237, PZ726225) show a close relationship with a human restvirus identified in China in 2019 (Song et al., 2024), supporting the hypothesis that restviruses are a respiratory-associated lineage. The third sequence (PZ726226) is related to a lineage of tombus-like viruses reported in individuals with lower respiratory tract infections in China with the highest similarity to a sequence from 2016 (OQ147340) (Ye et al., 2025). Due to an increase in serum IgG titers detected in patients negative for other respiratory viruses, this 2016 lineage was suggested to infect humans (Ye et al., 2025), however, further investigation is needed to determine the host of these viruses.

Lastly, we identified plant-infecting DNA and RNA viruses present in oral/nasal swabs of individuals in this study, likely due to recent consumption of plant material (Table 3, Supplementary Table 1). Plant-infecting DNA virus genomes identified here are members of the *Begomovirus* genus of the *Geminiviridae* family and include South African cassava mosaic virus (PZ726212-PZ726215), East African cassava mosaic Cameroon virus (PZ726165, PZ726166), and bean leaf curl Madagascar virus (PZ726164) (De Bruyn et al., 2016; Harimalala et al., 2012; Lefeuvre et al., 2007). Plant-infecting RNA virus genomes characterized here include a potexvirus in the *Alphaflexviridae* family (PZ726276) and tobacco mosaic viruses from the *Virgaviridae* family (PZ726223–PZ726224, PZ726227, PZ726229–PZ726230, PZ726234–PZ726236, PZ726244, PZ726251–PZ726252, PZ726257, PZ726261, and PZ726275). This detection of diverse plant viruses demonstrates the utility of human oral/nasal swabs for surveying viruses impacting key crops in the region such as cassava.

## Conclusions

The viral metagenomic workflow used here resulted in the identification of 650 coding-complete genomes across five human-infecting DNA virus families, seven human-infecting RNA families, one protist-infecting virus family, and three plant-infecting virus families. Human-infecting DNA and RNA viruses in Malagasy populations were generally closely related to virus lineages circulating globally and presented multiple pathogens of interest for public health in Madagascar. While some virus groups exhibited more of a geographic signature with a close relationship to sequences from African countries (e.g. rubella virus, HAstV-3), others were most similar to sequences obtained from samples of various geographic regions collected at similar time points in 2024 (e.g. human coronavirus HKU1 and OC43, enterovirus C105). We identified four viruses with unknown hosts including madahom bibyvirus 1 in *Picornaviridae* likely with a mammalian host given its phylogenetic position and madahom restvirus 1 and 2 and madahom tblv virus 1 which were most closely related to sequences from human respiratory samples.

In twenty-two individuals we detected at least one pathogenic DNA virus and in 61 individuals at least one pathogenic RNA virus. Multiple co-occurrences were documented (e.g., one individual was infected with viruses across four families). We did not see an association between malaria status and pathogenic virus detection, however, age was a significant driver of virus presence, with younger individuals having higher odds of pathogenic virus infection. We recognize that the absence of an observed relationship between *P. falciparum* and pathogenic virus infection may reflect the focus of our study on samples from the oral and nasal cavities which do not fully capture *P. falciparum’s* systemic capacity. The impact of age on virus infection may be driven by differences in acquired immunity with children more susceptible to pathogens in comparison with older individuals who may have acquired natural immunity to said pathogens. As children in the region seem to experience the heaviest burden of active protozoan and viral pathogen infections in conjunction with widespread malnutrition and anemia in Madagascar, community-led efforts in these villages supported by Health In Harmony are essential for providing resources such as sustainable protein sources (e.g., chicken farming, insect farming, aquaculture), agroforestry, clean water, and vaccinations.

Further, our work demonstrates the utility of oral swabs for understanding viruses present in important village crops (e.g. South African cassava mosaic virus, East African cassava mosaic Cameroon virus, bean leaf curl Madagascar virus). Our largest virus dataset was composed of 530 complete anellovirus genomes positioned across all nine of the established human-infecting anellovirus genera. While anelloviruses demonstrated high intra-individual variation, betatorqueviruses dominated across all age groups, highlighting a contrast with the domination of alphatorqueviruses in most human plasma studies. Further, in an anellovirus co-occurrence network, males and individuals with *P. falciparum* or pathogenic virus infections tended to share more anelloviruses, showing a putative connection between anelloviruses and host immune status. Broadly, this study exemplifies the value of working with small healthcare clinics in more rural, remote regions experiencing a high burden of infectious disease to build a landscape of actively circulating and co-infecting viral diversity in the area and to construct a foundation of high-quality DNA and RNA virus genomes for countries with limited viral genomic resources.

## Acknowledgments

We thank the communities in 29 villages within and near the Manombo Special Reserve (MSR) for their willingness to participate and provide samples for the research described here and their communities’ involvement in the research. We would like to thank the non-governmental organization Health In Harmony-Fahasalamana Mirindra, especially the incredible mobile healthcare clinic team in Farafangana who performed all sample and metadata collection along with Health In Harmony team members including Nina Finley and Laurent Raveloson who assisted with obtaining permits and following ethics protocols. We thank Zoo New England for their ongoing support and commitment to One Health research at the MSR. We thank the Comité Malgache d’Éthique pour les Sciences et les Technologies (CMEST), Duke University IRB, and Arizona State University IRB for research permissions.

## Human sampling

The human sampling described here was approved through the Comité Malgache d’Éthique pour les Sciences et les Technologies (CMEST) under permit N°194-2024 /AM/CMEST-P/ in Madagascar, the Duke University Campus Institutional Review Board under protocol #2025-0019, and the Arizona State University Institutional Review Board under study ID #STUDY00022785. Informed consent was obtained from all participants.

## Data availability

The virus sequences described in this study have been deposited in GenBank under accession numbers PZ726164–PZ726821 and PZ716971–PZ716984. Raw read sequence data for samples have been deposited under BioProject number PRJNA1290322, BioSample numbers SAMN61476770–SAMN61477009 and SRA numbers SRR39521765–SRR39522244.

## Table Captions

**Supplementary Table 1.** Summary of sequences characterized in this study and accompanying mean coverage and read count data.

**Supplementary Table 2.** Summary of sequences in the *Anelloviridae* family characterized in this study and accompanying mean coverage and read count data.

---

**Supplementary Figure 1.**
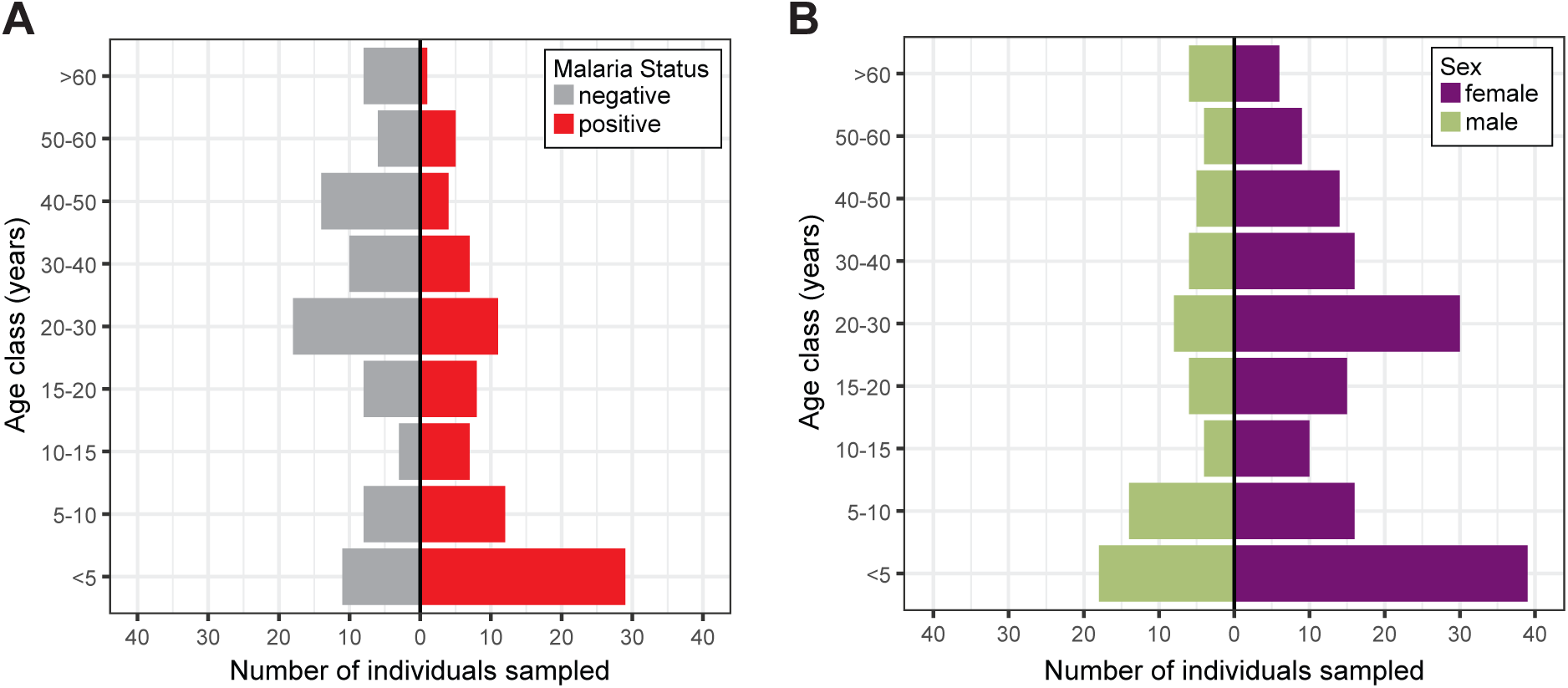
**(A)** Summary of 170 samples collected in this study for which malaria status metadata was available, stratified by age and malaria status. **(B)** Summary of 226 samples collected in this study for which demographic metadata was available, stratified by age and sex.

**Supplementary Figure 2.**
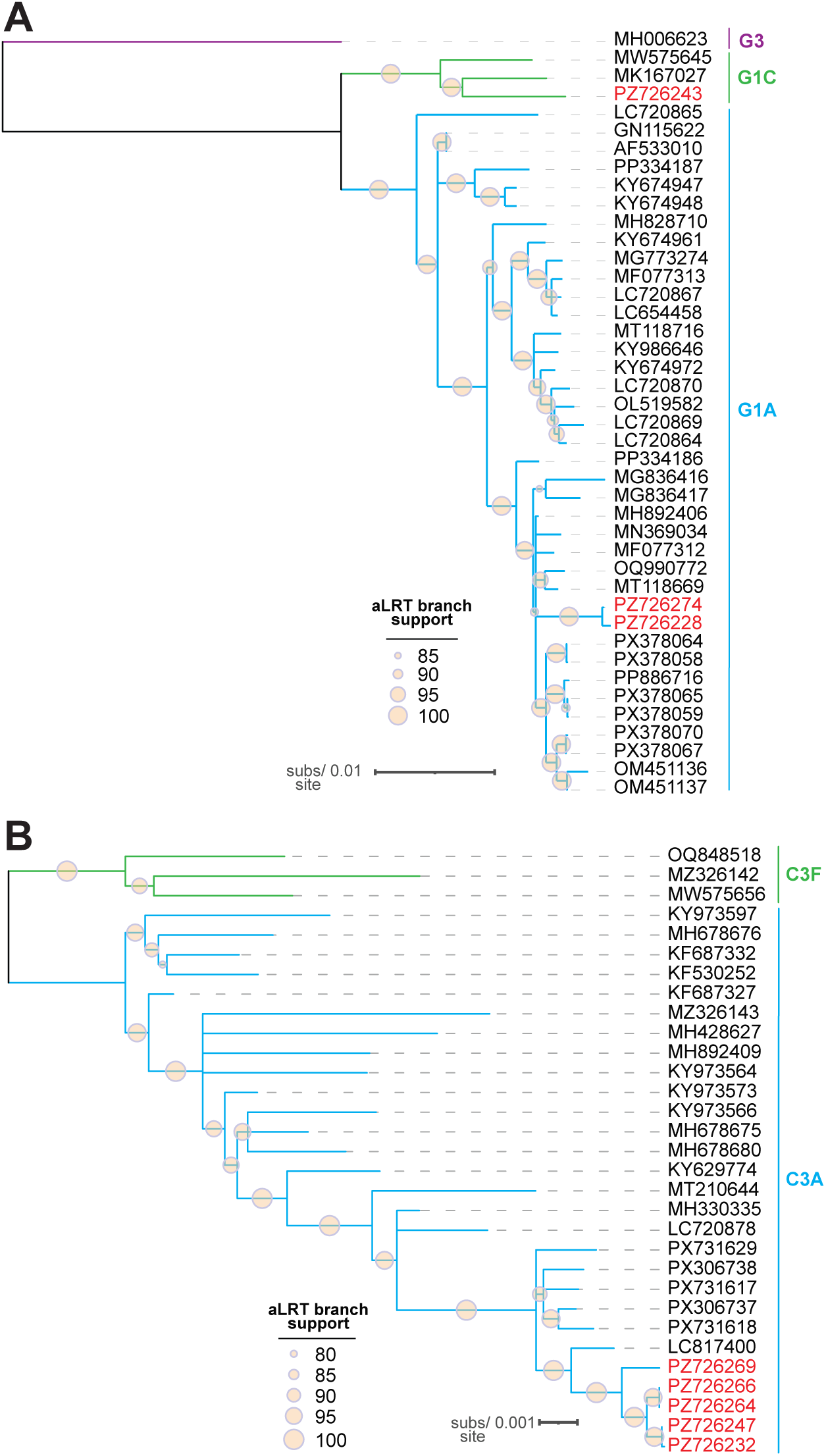
**(A)** Maximum likelihood phylogenetic tree of whole genome sequences of human parainfluenza virus type 2 G1A and G1C sequences rooted with G3 sequences. **(B)** Maximum likelihood phylogenetic tree of whole genome sequences of human parainfluenza virus type 3 C3A sequences rooted with C3F sequences. Viruses characterized in this study are shown in red font.

**Supplementary Figure 3.**
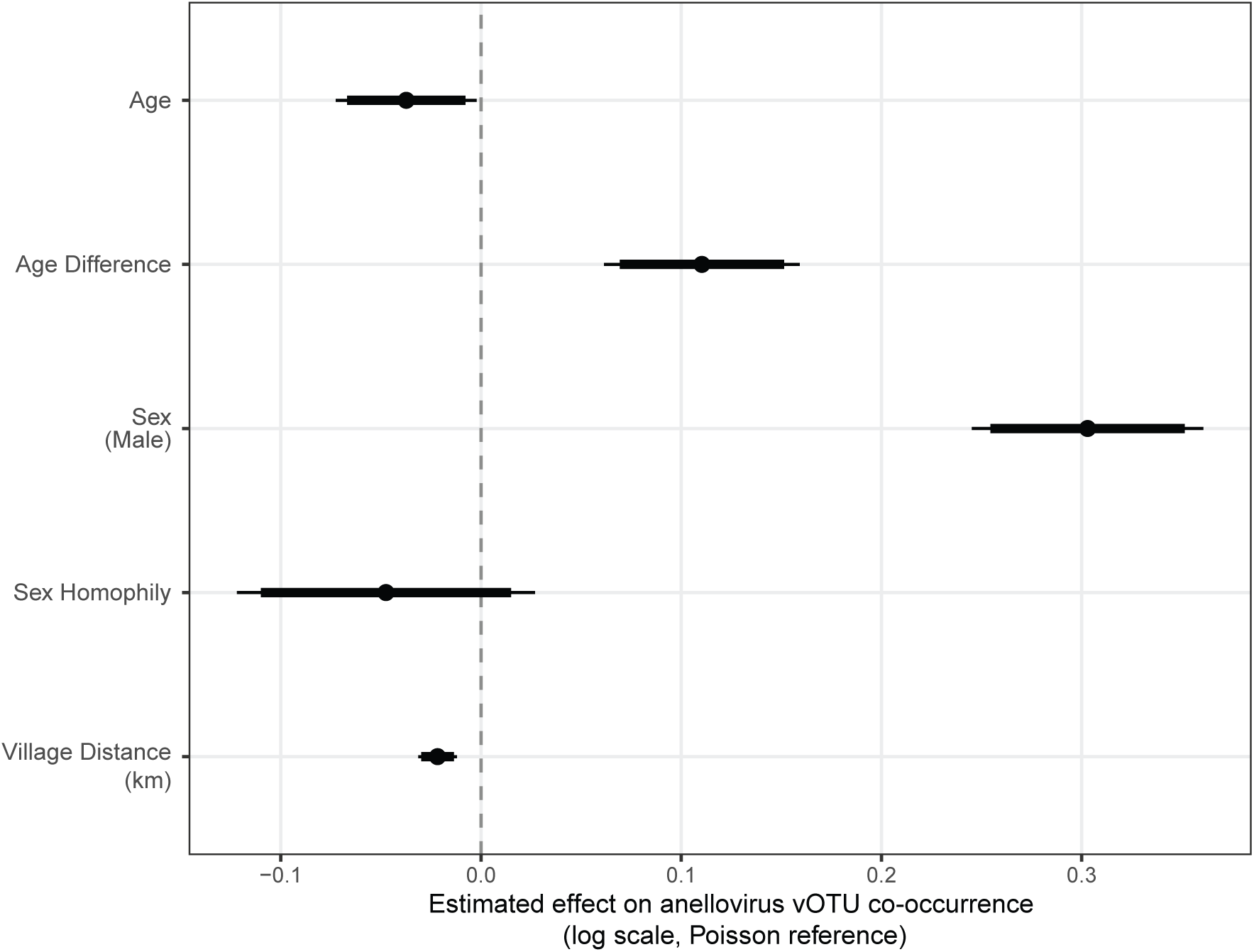
Using the broader dataset of 226 individuals for which demographic data was available, ERGM coefficient plot showing host traits associated with virus sharing. Note that the plot excludes the two network structural terms included in the model: “sum” and “nonzero”. Effect sizes are on the log-linear scale (i.e., for every 1 unit increase in a given variable, the number of shared viruses multiplies by e^b, where b is the estimated effect size; because the model includes a “nonzero” term, this rate does not directly represent the number of shared viruses). Positive coefficients represent that a given variable is associated with more shared viruses.

**Supplementary Figure 4.**
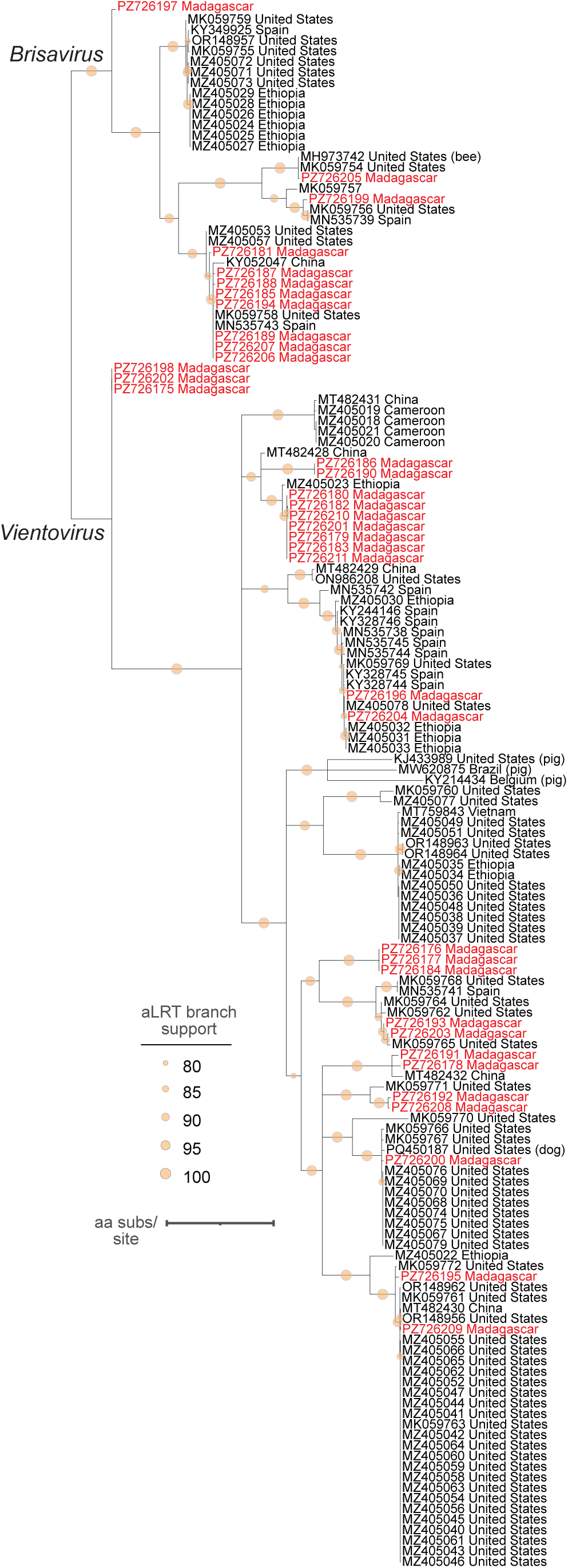
Maximum likelihood phylogenetic tree of vientovirus and brisavirus Rep amino acid sequences. Viruses characterized in this study are shown in red font.

## References

Abbas, A., Taylor, L.J., Collman, R.G., Bushman, F.D., 2021. ICTV Virus Taxonomy Profile: Redondoviridae. Journal of General Virology 102.

Abbas, A.A., Taylor, L.J., Dothard, M.I., Leiby, J.S., Fitzgerald, A.S., Khatib, L.A., Collman, R.G., Bushman, F.D., 2019. Redondoviridae, a Family of Small, Circular DNA Viruses of the Human Oro-Respiratory Tract Associated with Periodontitis and Critical Illness. Cell Host & Microbe 25, 719–729.e714.

Adiku, T., Dove, W., Grosjean, P., Combe, P., Nakagomi, T., Nakagomi, O., Hart, C., Cunliffe, N., 2010. Molecular characterization of rotavirus strains circulating among children with acute gastroenteritis in Madagascar during 2004–2005. Journal of Infectious Diseases 202, S175–S179.

Akello, J.O., Kamgang, R., Barbani, M.T., Suter-Riniker, F., Leib, S.L., Ramette, A., 2020. Epidemiology of human adenoviruses: a 20-year retrospective observational study in hospitalized patients in Bern, Switzerland. Clinical epidemiology, 353–366.

Allibert, C., 2008. Austronesian migration and the establishment of the Malagasy civilization: contrasted readings in linguistics, archaeology, genetics and cultural anthropology. Diogenes 55, 7–16.

Anfruns-Estrada, E., Sabrià, A., Fuentes, C., Sabaté, S., Razquin, E., Cornejo, T., Bartolomé, R., Torner, N., Izquierdo, C., Soldevila, N., 2020. Detection of norovirus in saliva samples from acute gastroenteritis cases and asymptomatic subjects: association with age and higher shedding in stool. Viruses 12, 1369.

Aroney, S.T.N., Newell, R.J.P., Nissen, J.N., Camargo, A.P., Tyson, G.W., Woodcroft, B.J., 2025. CoverM: read alignment statistics for metagenomics. Bioinformatics 41, btaf147.

Babakir-Mina, M., Ciccozzi, M., Alteri, C., Polchi, P., Picardi, A., Greco, F., Lucarelli, G., Arcese, W., Perno, C.F., Ciotti, M., 2009. Excretion of the novel polyomaviruses KI and WU in the stool of patients with hematological disorders. Journal of Medical Virology 81, 1668–1673.

Baele, G., Ji, X., Hassler, G.W., McCrone, J.T., Shao, Y., Zhang, Z., Holbrook, A.J., Lemey, P., Drummond, A.J., Rambaut, A., Suchard, M.A., 2025. BEAST X for Bayesian phylogenetic, phylogeographic and phylodynamic inference. Nature Methods 22, 1653–1656.

Bates, D., Mächler, M., Bolker, B., Walker, S., 2015. Fitting linear mixed-effects models using lme4. Journal of statistical software 67, 1–48.

Bian, L., Wang, Y., Yao, X., Mao, Q., Xu, M., Liang, Z., 2015. Coxsackievirus A6: a new emerging pathogen causing hand, foot and mouth disease outbreaks worldwide. Expert review of anti-infective therapy 13, 1061–1071.

Bigoni, J., Catarino, R., Benski, C., Viviano, M., Munoz, M., Tilahizandry, H., Petignat, P., Vassilakos, P., 2018. High burden of human papillomavirus infection in Madagascar: comparison with other sexually transmitted infections. Infectious Diseases: Research and Treatment 11, 1178633717752686.

Bird, B.H., Khristova, M.L., Rollin, P.E., Ksiazek, T.G., Nichol, S.T., 2007. Complete genome analysis of 33 ecologically and biologically diverse Rift Valley fever virus strains reveals widespread virus movement and low genetic diversity due to recent common ancestry. J Virol 81, 2805–2816.

Bolger, A.M., Lohse, M., Usadel, B., 2014. Trimmomatic: a flexible trimmer for Illumina sequence data. Bioinformatics 30, 2114–2120.

Boros, Á., Orlovácz, K., Pankovics, P., Szekeres, S., Földvári, G., Fahsbender, E., Delwart, E., Reuter, G., 2019. Diverse picornaviruses are prevalent among free-living and laboratory rats (Rattus norvegicus) in Hungary and can cause disseminated infections. Infection, Genetics and Evolution 75, 103988.

Brister, J.R., Ako-Adjei, D., Bao, Y., Blinkova, O., 2015. NCBI viral genomes resource. Nucleic Acids Res 43, D571–577.

Bryce, J., Boschi-Pinto, C., Shibuya, K., Black, R.E., 2005. WHO estimates of the causes of death in children. Lancet 365, 1147–1152.

Buchfink, B., Xie, C., Huson, D.H., 2015. Fast and sensitive protein alignment using DIAMOND. Nature Methods 12, 59–60.

Carroll, S.A., Reynes, J.M., Khristova, M.L., Andriamandimby, S.F., Rollin, P.E., Nichol, S.T., 2011. Genetic evidence for Rift Valley fever outbreaks in Madagascar resulting from virus introductions from the East African mainland rather than enzootic maintenance. J Virol 85, 6162–6167.

Cordey, S., Laubscher, F., Hartley, M.A., Junier, T., Keitel, K., Docquier, M., Guex, N., Iseli, C., Vieille, G., Le Mercier, P., Gleizes, A., Samaka, J., Mlaganile, T., Kagoro, F., Masimba, J., Said, Z., Temba, H., Elbanna, G.H., Tapparel, C., Zanella, M.C., Xenarios, I., Fellay, J., D’Acremont, V., Kaiser, L., 2021. Blood virosphere in febrile Tanzanian children. Emerg Microbes Infect 10, 982–993.

Cui, L., Wu, B., Zhu, X., Guo, X., Ge, Y., Zhao, K., Qi, X., Shi, Z., Zhu, F., Sun, L., Zhou, M., 2017. Identification and genetic characterization of a novel circular single-stranded DNA virus in a human upper respiratory tract sample. Arch Virol 162, 3305–3312.

Cui, S., Guo, R., Chen, C., Zhang, Y., Meng, J., Liu, L., Li, Y., Kang, Z., Li, S., Yan, Q., 2024. Next-Generation Sequencing for Characterizing Respiratory Tract Virome and Improving Detection of Viral Pathogens in Children With Pneumonia. Influenza and Other Respiratory Viruses 18, e13362.

Daleno, C., Piralla, A., Scala, A., Baldanti, F., Usonis, V., Principi, N., Esposito, S., 2012. Complete genome sequence of a novel human enterovirus C (HEV-C117) identified in a child with community-acquired pneumonia. American Society for Microbiology 1752 N St., NW, Washington, DC.

De Bruyn, A., Harimalala, M., Zinga, I., Mabvakure, B.M., Hoareau, M., Ravigné, V., Walters, M., Reynaud, B., Varsani, A., Harkins, G.W., 2016. Divergent evolutionary and epidemiological dynamics of cassava mosaic geminiviruses in Madagascar. BMC evolutionary biology 16, 182.

De Vlaminck, I., Khush, K.K., Strehl, C., Kohli, B., Luikart, H., Neff, N.F., Okamoto, J., Snyder, T.M., Cornfield, D.N., Nicolls, M.R., Weill, D., Bernstein, D., Valantine, H.A., Quake, S.R., 2013. Temporal response of the human virome to immunosuppression and antiviral therapy. Cell 155, 1178–1187.

Dewar, R.E., Richard, A.F., 2012. Madagascar: A History of Arrivals, What Happened, and Will Happen Next*. Annual Review of Anthropology 41, 495–517.

Dhingra, A., Hage, E., Ganzenmueller, T., Böttcher, S., Hofmann, J., Hamprecht, K., Obermeier, P., Rath, B., Hausmann, F., Dobner, T., Heim, A., 2019. Molecular Evolution of Human Adenovirus (HAdV) Species C. Scientific Reports 9, 1039.

Drummond, A.J., Ho, S.Y.W., Phillips, M.J., Rambaut, A., 2006. Relaxed phylogenetics and dating with confidence. PLoS biology 4, e88.

Du, J., Zhu, T., Zhuang, L., Zhang, P.H., Zhang, X.A., Lu, Q.B., Liu, W., 2019. Identification and complete genome characterization of human enterovirus 117 from a child with pneumonia in China. Arch Virol 164, 1489–1492.

Fall, A., Zhuang, T.X., Dodge, A., Abdullah, O., Norton, J.M., Villafuerte, D., Pekosz, A., Klein, E., Mostafa, H.H., 2025. Molecular Characterization of Emerging and Uncommon Enteroviruses C104, C105, and C109 in Respiratory Samples from Maryland, USA, 2018–2024. Viruses 17, 1183.

Fazeli, M.M., Heydari Sirat, S., Shatizadeh Malekshahi, S., 2025. Novel Human Polyomaviruses Discovered From 2007 to the Present: An Update of Current Knowledge. Reviews in Medical Virology 35, e70017.

Fernandez-Garcia, M.D., Camacho, J., Diez-Fuertes, F., Ruiz de Pedro, E., García-Ibañez, N., Navascués, A., Berengua, C., Antequera-Rodriguez, P., Ruiz-García, M., Pastor-Fajardo, M.T., Cabrerizo, M., 2025. Detections of rare enterovirus C105 linked to an emerging novel clade, Spain, 2019 to 2024. Euro Surveill 30.

Freeman, M.C., Garn, J.V., Sclar, G.D., Boisson, S., Medlicott, K., Alexander, K.T., Penakalapati, G., Anderson, D., Mahtani, A.G., Grimes, J.E.T., Rehfuess, E.A., Clasen, T.F., 2017. The impact of sanitation on infectious disease and nutritional status: A systematic review and meta-analysis. International Journal of Hygiene and Environmental Health 220, 928–949.

Fu, L., Niu, B., Zhu, Z., Wu, S., Li, W., 2012. CD-HIT: accelerated for clustering the next-generation sequencing data. Bioinformatics 28, 3150–3152.

Fulkerson, H.L., Nogalski, M.T., Collins-McMillen, D., Yurochko, A.D., 2021. Overview of human cytomegalovirus pathogenesis. Human cytomegaloviruses: methods and protocols, 1–18.

Garnett, C., Erdman, D., Xu, W., Gooding, L.R., 2002. Prevalence and quantitation of species C adenovirus DNA in human mucosal lymphocytes. Journal of virology 76, 10608–10616.

Garsevanidze, E., Kislaya, I., Kutz, J.-M., Rasamoelina, T., Ratefiarisoa, S., Razafindrakoto, R., Rasolojaona, Z.T., Razafindralava, N.M., Totofotsy, O., Zafinimampera, A.O., Randrianarisoa, T., Lassmann, M., Remkes, A., Brito, A., Andrianarimanana, D.K., Rakotomalala, R.S., Rakotomalala, Z., Rausche, P., Hey, J., McKay-Chopin, S., May, J., Hampl, M., Marchese, V., Gheit, T., Rakotoarivelo, R.A., Fusco, D., 2026. Circulating genotypes of human papillomavirus in adult women of reproductive age from the Boeny region of Madagascar: a cross-sectional study to explore needs and opportunities for HPV vaccination in the country. BMC Infectious Diseases 26, 425.

Gatherer, D., Depledge, D.P., Hartley, C.A., Szpara, M.L., Vaz, P.K., Benkő, M., Brandt, C.R., Bryant, N.A., Dastjerdi, A., Doszpoly, A., 2021. ICTV virus taxonomy profile: Herpesviridae 2021. Journal of General Virology 102, 001673.

Graf, E.H., Simmon, K.E., Tardif, K.D., Hymas, W., Flygare, S., Eilbeck, K., Yandell, M., Schlaberg, R., 2016. Unbiased detection of respiratory viruses by use of RNA sequencing-based metagenomics: a systematic comparison to a commercial PCR panel. Journal of clinical microbiology 54, 1000–1007.

Gregg, N.M., 1991. Congenital cataract following German measles in the mother. Epidemiology & Infection 107, iii–xiv.

Griffin, D.E., 2022. Why does viral RNA sometimes persist after recovery from acute infections? PLoS biology 20, e3001687.

Grinde, B., 2013. Herpesviruses: latency and reactivation–viral strategies and host response. Journal of oral microbiology 5, 22766.

Guillebaud, J., Bernardson, B., Randriambolamanantsoa, T.H., Randrianasolo, L., Randriamampionona, J.L., Marino, C.A., Rasolofo, V., Randrianarivelojosia, M., Vigan-Womas, I., Stivaktas, V., 2018. Study on causes of fever in primary healthcare center uncovers pathogens of public health concern in Madagascar. PLoS neglected tropical diseases 12, e0006642.

Harimalala, M., Lefeuvre, P., De Bruyn, A., Tiendrébéogo, F., Hoareau, M., Villemot, J., Ranomenjanahary, S., Andrianjaka, A., Reynaud, B., Lett, J.-M., 2012. A novel cassava-infecting begomovirus from Madagascar: cassava mosaic Madagascar virus. Archives of virology 157, 2027–2030.

Hause, B.M., Nelson, E., Christopher-Hennings, J., 2021. Identification of a novel statovirus in a faecal sample from a calf with enteric disease. Journal of General Virology 102, 001655.

Hijmans, R.J., Williams, E., Vennes, C., Hijmans, M.R.J., 2017. Package ‘geosphere’. Spherical trigonometry 1, 1–45.

Hill, V., Baele, G., 2019. Bayesian estimation of past population dynamics in BEAST 1.10 using the Skygrid coalescent model. Molecular biology and evolution 36, 2620–2628.

Hirose, E., Minagawa, H., Adachi, H., Suzuki, M., Nakamura, N., Saito, N., Ito, M., Mitsuoka, C., Sato, K., Yasui, Y., 2025. Detection of Recombinant Type of Human Adenovirus C Composed of Types 89 and 5 in Aichi Prefecture, Japan (2016–2019). Japanese Journal of Infectious Diseases 78, 139–142.

Hoffmann, J., Rabezanahary, H., Randriamarotia, M., Ratsimbasoa, A., Najjar, J., Vernet, G., Contamin, B., Paranhos-Baccala, G., 2012. Viral and atypical bacterial etiology of acute respiratory infections in children under 5 years old living in a rural tropical area of Madagascar.

Horner, L.M., Poulter, M.D., Brenton, J.N., Turner, R.B., 2015. Acute Flaccid Paralysis Associated with Novel Enterovirus C105. Emerg Infect Dis 21, 1858–1860.

Howard, C.T., de Pee, S., Sari, M., Bloem, M.W., Semba, R.D., 2007. Association of diarrhea with anemia among children under age five living in rural areas of Indonesia. J Trop Pediatr 53, 238–244.

Hunter, D.R., Handcock, M.S., Butts, C.T., Goodreau, S.M., Morris, M., 2008. ergm: A package to fit, simulate and diagnose exponential-family models for networks. Journal of statistical software 24, 1–29.

Janowski, A.B., Krishnamurthy, S.R., Lim, E.S., Zhao, G., Brenchley, J.M., Barouch, D.H., Thakwalakwa, C., Manary, M.J., Holtz, L.R., Wang, D., 2017. Statoviruses, a novel taxon of RNA viruses present in the gastrointestinal tracts of diverse mammals. Virology 504, 36–44.

Kaczorowska, J., Deijs, M., Klein, M., Bakker, M., Jebbink, M.F., Sparreboom, M., Kinsella, C.M., Timmerman, A.L., van der Hoek, L., 2022. Diversity and Long-Term Dynamics of Human Blood Anelloviruses. J Virol 96, e0010922.

Kaczorowska, J., Van Der Hoek, L., 2020. Human anelloviruses: diverse, omnipresent and commensal members of the virome. FEMS Microbiology Reviews 44, 305–313.

Kalyaanamoorthy, S., Minh, B.Q., Wong, T.K., Von Haeseler, A., Jermiin, L.S., 2017. ModelFinder: fast model selection for accurate phylogenetic estimates. Nature methods 14, 587–589.

Katoh, K., Standley, D.M., 2013. MAFFT multiple sequence alignment software version 7: improvements in performance and usability. Mol Biol Evol 30, 772–780.

Keeler, E.L., Merenstein, C., Reddy, S., Taylor, L.J., Cobián-Güemes, A.G., Zankharia, U., Collman, R.G., Bushman, F.D., 2023. Widespread, human-associated redondoviruses infect the commensal protozoan Entamoeba gingivalis. Cell host & microbe 31, 58–68. e55.

Kenmoe, S., Sadeuh-Mba, S.A., Vernet, M.A., Penlap Beng, V., Vabret, A., Njouom, R., 2021. Molecular epidemiology of Enteroviruses and Rhinoviruses in patients with acute respiratory infections in Yaounde, Cameroon. Influenza Other Respir Viruses 15, 641–650.

Kennedy, J.L., Kincaid, J.C., Schwalm, K.C., Stoner, A.N., Abramo, T.J., Thompson, T.M., Hardin, O., Putt, C., Dinwiddie, D.L., 2017. Genome Sequences of Three Novel Isolates of Human Parainfluenza Virus 2 Associated with Acute Respiratory Infection. Genome Announcements 5, 10.1128/genomea.00784-00717.

Khanal, S., Ghimire, P., Dhamoon, A.S., 2018. The repertoire of adenovirus in human disease: the innocuous to the deadly. Biomedicines 6, 30.

Kinsella, C.M., Deijs, M., Becker, C., Broekhuizen, P., van Gool, T., Bart, A., Schaefer, A.S., van der Hoek, L., 2022. Host prediction for disease-associated gastrointestinal cressdnaviruses. Virus Evolution.

Kleinschmidt-DeMasters, B.K., Gilden, D.H., 2001. The Expanding Spectrum of Herpesvirus Infections of the Nervous System. Brain Pathology 11, 440–451.

Kraberger, S., Cook, C.N., Schmidlin, K., Fontenele, R.S., Bautista, J., Smith, B., Varsani, A., 2019. Diverse single-stranded DNA viruses associated with honey bees (Apis mellifera). Infection, Genetics and Evolution 71, 179–188.

Krivitsky, P.N., 2012. Exponential-family random graph models for valued networks. Electronic journal of statistics 6, 1100.

Kutz, J.-M., Rausche, P., Rasamoelina, T., Ratefiarisoa, S., Razafindrakoto, R., Klein, P., Jaeger, A., Rakotomalala, R.S., Rakotomalala, Z., Randrianasolo, B.S., McKay-Chopin, S., May, J., Rakotozandrindrainy, R., Puradiredja, D.I., Sicuri, E., Hampl, M., Lorenz, E., Gheit, T., Rakotoarivelo, R.A., Fusco, D., 2023. Female genital schistosomiasis, human papilloma virus infection, and cervical cancer in rural Madagascar: a cross sectional study. Infectious Diseases of Poverty 12, 89.

Lambert, N., Strebel, P., Orenstein, W., Icenogle, J., Poland, G.A., 2015. Rubella. The Lancet 385, 2297–2307.

Lambisia, A.W., Mutunga, M., Katama, E.N., Agoti, C.N., Houldcroft, C.J., 2024. Multispecies Cocirculation of Adenoviruses Identified by Next-Generation Sequencing During an Acute Gastroenteritis Outbreak in Coastal Kenya in 2023. Open Forum Infect Dis 11, ofae505.

Lau, S.K.P., Woo, P.C.Y., Li, K.S.M., Zhang, H.-J., Fan, R.Y.Y., Zhang, A.J.X., Chan, B.C.C., Lam, C.S.F., Yip, C.C.Y., Yuen, M.-C., Chan, K.-H., Chen, Z.-W., Yuen, K.-Y., 2016. Identification of Novel Rosavirus Species That Infects Diverse Rodent Species and Causes Multisystemic Dissemination in Mouse Model. PLOS Pathogens 12, e1005911.

Lee, B., Damon, C.F., Platts-Mills, J.A., 2020. Pediatric acute gastroenteritis associated with adenovirus 40/41 in low-income and middle-income countries. Current opinion in infectious diseases 33, 398–403.

Lefeuvre, P., Martin, D.P., Hoareau, M., Naze, F., Delatte, H., Thierry, M., Varsani, A., Becker, N., Reynaud, B., Lett, J.-M., 2007. Begomovirus ‘melting pot’in the south-west Indian Ocean islands: molecular diversity and evolution through recombination. Journal of General Virology 88, 3458–3468.

Letunic, I., Bork, P., 2021. Interactive Tree Of Life (iTOL) v5: an online tool for phylogenetic tree display and annotation. Nucleic Acids Res 49, W293–w296.

Li, D., Liu, C.-M., Luo, R., Sadakane, K., Lam, T.-W., 2015. MEGAHIT: an ultra-fast single-node solution for large and complex metagenomics assembly via succinct de Bruijn graph. Bioinformatics 31, 1674–1676.

Li, M., Zhang, T., Gong, C., Li, A., Luo, M., Dong, M., Huang, F., 2019. Human Enterovirus C105, China, 2017. Emerg Infect Dis 25, 1414–1416.

Li, Y., Cao, L., Ye, M., Xu, R., Chen, X., Ma, Y., Tian, R.-R., Liu, F.-L., Zhang, P., Kuang, Y.-Q., Zheng, Y.-T., Zhang, C., 2022. Plasma Virome Reveals Blooms and Transmission of Anellovirus in Intravenous Drug Users with HIV-1, HCV, and/or HBV Infections. Microbiology Spectrum 10, e01447–01422.

Lim, E.S., Cao, S., Holtz, L.R., Antonio, M., Stine, O.C., Wang, D., 2014. Discovery of rosavirus 2, a novel variant of a rodent-associated picornavirus, in children from The Gambia. Virology 454–455, 25-33.

Liu, D.X., Liang, J.Q., Fung, T.S., 2021. Human coronavirus-229E,-OC43,-NL63, and-HKU1 (Coronaviridae). Encyclopedia of virology, 428.

Lockwood, S., Ranaivoson, H.C., Randriambolamanantsoa, T.H., Razanajatovo, N., Raharinosy, V., Ahyong, V., Héraud, J.-M., Dussart, P., Lacoste, V., Brook, C.E., 2026. Identifying viral infections through metagenomic Next Generation Sequencing of undiagnosed respiratory fevers in Madagascar (2014–2019). BMC Infectious Diseases.

Lukashev, A.N., Drexler, J.F., Kotova, V.O., Amjaga, E.N., Reznik, V.I., Gmyl, A.P., Grard, G., Taty Taty, R., Trotsenko, O.E., Leroy, E.M., 2012. Novel serotypes 105 and 116 are members of distinct subgroups of human enterovirus C. Journal of general virology 93, 2357–2362.

Maeder, M.N., Rabezanahary, H.M., Zafindraibe, N.J., Randriatiana, M.R., Rasamoelina, T., Rakotoarivo, A.T., Vanhems, P., Hoffmann, J., Bénet, T., Rakoto Andrianarivelo, M., 2016. Sickle-cell disease in febrile children living in a rural village of Madagascar and association with malaria and respiratory infections. BMC hematology 16, 30.

Majumdar, M., Celma, C., Pegg, E., Polra, K., Dunning, J., Martin, J., 2021. Detection and Typing of Human Enteroviruses from Clinical Samples by Entire-Capsid Next Generation Sequencing. Viruses 13, 641.

Mansour, O., Fadeev, A.V., Perederiy, A.A., Zadirienko, M.I., Danilenko, D.M., Lioznov, D.A., Komissarov, A.B., 2025. Genetic Diversity and Molecular Analysis of Human Parainfluenza Virus Type 3 in Saint Petersburg (Russia) in 2017-2023: Emergence of a New Phylogenetic Cluster. Viruses 17.

Mashhadi Abolghasem Shirazi, M., 2026. Comparative innate immune responses across major RNA and DNA viral infections: Mechanisms, immunopathology, and therapeutic perspectives. Hum Vaccin Immunother 22, 2714657.

Mawson, A.R., Majumdar, S., 2017. Malaria, Epstein–Barr virus infection and the pathogenesis of Burkitt’s lymphoma. International Journal of Cancer 141, 1849–1855.

McElvania TeKippe, E., Wylie, K.M., Deych, E., Sodergren, E., Weinstock, G., Storch, G.A., 2012. Increased prevalence of anellovirus in pediatric patients with fever. PloS one 7, e50937.

Miller, E., Cradock-Watson, J., Pollock, T., 1982. Consequences of confirmed maternal rubella at successive stages of pregnancy. The Lancet 320, 781–784.

Minh, B.Q., Schmidt, H.A., Chernomor, O., Schrempf, D., Woodhams, M.D., von Haeseler, A., Lanfear, R., 2020. IQ-TREE 2: New Models and Efficient Methods for Phylogenetic Inference in the Genomic Era. Mol Biol Evol 37, 1530–1534.

Mokoena, F., Esona, M.D., Seheri, L.M., Nyaga, M.M., Magagula, N.B., Mukaratirwa, A., Mulindwa, A., Abebe, A., Boula, A., Tsolenyanu, E., 2021. Whole genome analysis of African G12P [6] and G12P [8] Rotaviruses provides evidence of porcine-human reassortment at NSP2, NSP3, and NSP4. Frontiers in Microbiology 11, 604444.

Moormann, A.M., Bailey, J.A., 2016. Malaria - how this parasitic infection aids and abets EBV-associated Burkitt lymphomagenesis. Curr Opin Virol 20, 78–84.

Moyen, N., Thiberville, S.D., Pastorino, B., Nougairede, A., Thirion, L., Mombouli, J.V., Dimi, Y., Leparc-Goffart, I., Capobianchi, M.R., Lepfoundzou, A.D., de Lamballerie, X., 2014. First reported chikungunya fever outbreak in the republic of Congo, 2011. PLoS One 9, e115938.

Mueller, A., Simon, A., Gillen, J., Schildgen, V., Tillmann, R.L., Reiter, K., Schildgen, O., 2009. Polyomaviruses KI and WU in children with respiratory tract infection. Archives of virology 154, 1605–1608.

Muhire, B.M., Varsani, A., Martin, D.P., 2014. SDT: a virus classification tool based on pairwise sequence alignment and identity calculation. PLoS One 9, e108277.

Osman, A., 2016. Protein energy malnutrition and susceptibility to viral infections as zika and influenza viruses. J Nutr Food Sci 6, 2.

Paietta, E.N., Johnston, R.A., Kraberger, S., Randrianarisoa, S.F., Razanamahenina, T.T., Ramboninarimalala, A., Velontsara, J.-B., Raherinirina, T.G., Raveloson, L., Finley, N.L., 2026a. Mammal-infecting DNA viruses identified in lemurs and rodents in Madagascar mirror the evolutionary history of their hosts. Microbial Genomics 12, 001728.

Paietta, E.N., Johnston, R.A., Randrianarisoa, S.F., DeSisto, C.M.M., Kraberger, S., Martin, D., Razanamahenina, T.T., Ramboninarimalala, A., Velontsara, J.-B., Raherinirina, T.G., Rasendrinirina, V., Raveloson, L., Luo, Y., Finley, N.L., Baitchman, E., Yoder, A.D., Varsani, A., 2026b. Host–anellovirus interactions in an island ecosystem: non-human primates and rodents in Madagascar harbour diverse, rich anellovirus populations. Microbial Genomics 12.

Paietta, E.N., Lefkowitz, E.J., Van Der Pol, W.J., Hendrickson, R.C., Johnston, R.A., Randrianarisoa, S.F., Kraberger, S., Razanamahenina, T.T., Ramboninarimalala, A., Raherinirina, T.G., 2026c. Divergent poxvirus identified in a non-native black rat from Madagascar. Virology, 111021.

Papaventsis, D.C., Dove, W., Cunliffe, N.A., Nakagomi, O., Combe, P., Grosjean, P., Hart, C.A., 2008. Human astrovirus gastroenteritis in children, Madagascar, 2004-2005. Emerg Infect Dis 14, 844–846.

Pawełczyk, M., Kowalski, M.L., 2017. The Role of Human Parainfluenza Virus Infections in the Immunopathology of the Respiratory Tract. Current Allergy and Asthma Reports 17, 16.

Piralla, A., Daleno, C., Girello, A., Esposito, S., Baldanti, F., 2015. Circulation of two Enterovirus C105 (EV-C105) lineages in Europe and Africa. Journal of General Virology 96, 1374–1379.

Prendergast, A.J., Kelly, P., 2016. Interactions between intestinal pathogens, enteropathy and malnutrition in developing countries. Curr Opin Infect Dis 29, 229–236.

Prüss-Ustün, A., Bartram, J., Clasen, T., Colford Jr, J.M., Cumming, O., Curtis, V., Bonjour, S., Dangour, A.D., De France, J., Fewtrell, L., Freeman, M.C., Gordon, B., Hunter, P.R., Johnston, R.B., Mathers, C., Mäusezahl, D., Medlicott, K., Neira, M., Stocks, M., Wolf, J., Cairncross, S., 2014. Burden of disease from inadequate water, sanitation and hygiene in low- and middle-income settings: a retrospective analysis of data from 145 countries. Tropical Medicine & International Health 19, 894–905.

Rahajamanana, V.L., Raboba, J.L., Rakotozanany, A., Razafindraibe, N.J., Andriatahirintsoa, E.J.P.R., Razafindrakoto, A.C., Mioramalala, S.A., Razaiarimanga, C., Weldegebriel, G.G., Burnett, E., Mwenda, J.M., Seheri, M., Mphahlele, M.J., Robinson, A.L., 2018. Impact of rotavirus vaccine on all-cause diarrhea and rotavirus hospitalizations in Madagascar. Vaccine 36, 7198–7204.

Rajagopala, S.V., Bakhoum, N.G., Pakala, S.B., Shilts, M.H., Rosas-Salazar, C., Mai, A., Boone, H.H., McHenry, R., Yooseph, S., Halasa, N., 2021. Metatranscriptomics to characterize respiratory virome, microbiome, and host response directly from clinical samples. Cell reports methods 1.

Rajatonirina, S., Heraud, J.-M., Orelle, A., Randrianasolo, L., Razanajatovo, N., Rajaona, Y.R., Randrianarivo-Solofoniaina, A.E., Rakotomanana, F., Richard, V., 2012. The spread of influenza A (H1N1) pdm09 virus in Madagascar described by a sentinel surveillance network. PloS one 7, e37067.

Rajatonirina, S., Rakotosolofo, B., Rakotomanana, F., Randrianasolo, L., Ratsitoharina, M., Raharinandrasana, H., Heraud, J.-M., Richard, V., 2013a. Excess mortality associated with the 2009 A (H1N1) v influenza pandemic in Antananarivo, Madagascar. Epidemiology & Infection 141, 745–750.

Rajatonirina, S., Razanajatovo, N.H., Ratsima, E.H., Orelle, A., Ratovoson, R., Andrianirina, Z.Z., Andriatahina, T., Ramparany, L., Herindrainy, P., Randrianirina, F., 2013b. Outcome risk factors during respiratory infections in a paediatric ward in Antananarivo, Madagascar 2010–2012. PLoS One 8, e72839.

Rakau, K.G., Nyaga, M.M., Gededzha, M.P., Mwenda, J.M., Mphahlele, M.J., Seheri, L.M., Steele, A.D., 2021. Genetic characterization of G12P [6] and G12P [8] rotavirus strains collected in six African countries between 2010 and 2014. BMC Infectious Diseases 21, 107.

Rakoto-Andrianarivelo, M., Gumede, N., Jegouic, S., Balanant, J., Andriamamonjy, S.N., Rabemanantsoa, S., Birmingham, M., Randriamanalina, B., Nkolomoni, L., Venter, M., Schoub, B.D., Delpeyroux, F., Reynes, J.M., 2008. Reemergence of recombinant vaccine-derived poliovirus outbreak in Madagascar. J Infect Dis 197, 1427–1435.

Rakotomalala, F.A., Butel, C., Rasamoelina, T., Serrano, L., Vidal, N., Randriarimanana, S.H.D., Maharavo, L., Randriamananjara, H.N., Fernandez-Nuñez, N., Rabetokotany, F.R., 2024. High HIV-1 genetic diversity and low prevalence of transmitted drug resistance among treatment-naive people living with HIV in Madagascar. Infection, Genetics and Evolution 125, 105679.

Rambaut, A., Drummond, A.J., Xie, D., Baele, G., Suchard, M.A., 2018. Posterior summarization in Bayesian phylogenetics using Tracer 1.7. Systematic biology 67, 901–904.

Ramesh, A., Bailey, E.S., Ahyong, V., Langelier, C., Phelps, M., Neff, N., Sit, R., Tato, C., DeRisi, J.L., Greer, A.G., 2021. Metagenomic characterization of swine slurry in a North American swine farm operation. Scientific Reports 11, 16994.

Randremanana, R., Randrianirina, F., Gousseff, M., Dubois, N., Razafindratsimandresy, R., Hariniana, E.R., Garin, B., Randriamanantena, A., Rakotonirina, H.C., Ramparany, L., Ramarokoto, C.E., Rakotomanana, F., Ratsitorahina, M., Rajatonirina, S., Talarmin, A., Richard, V., 2012. Case-Control Study of the Etiology of Infant Diarrheal Disease in 14 Districts in Madagascar. PLoS ONE 7, e44533.

Randremanana, R.V., Razafindratsimandresy, R., Andriatahina, T., Randriamanantena, A., Ravelomanana, L., Randrianirina, F., Richard, V., 2016. Etiologies, risk factors and impact of severe diarrhea in the under-fives in Moramanga and Antananarivo, Madagascar. PLoS One 11, e0158862.

Randriambolamanantsoa, T.H., Razanajatovo, N.H., Ranaivoson, H.C., Randrianasolo, L., Rabarison, H.J., Razafimanjato, H., Ratsimbazafy, A., Rakoto, D.A.D., Heraud, J.-M., Lacoste, V., 2024. Identifying climatic drivers of respiratory syncytial virus (RSV) seasonality in Antananarivo, Madagascar, 2011–2021: a sentinel surveillance study. BMJ Public Health 2.

Razafindratsimandresy, R., Heraud, J.M., Ramarokoto, C.E., Rabemanantsoa, S., Randremanana, R., Andriamamonjy, N.S., Richard, V., Reynes, J.M., 2013. Rotavirus genotypes in children in the community with diarrhea in Madagascar. J Med Virol 85, 1652–1660.

Razafindratsimandresy, R., Joffret, M.L., Andriamandimby, S.F., Andriamamonjy, S., Rabemanantsoa, S., Richard, V., Delpeyroux, F., Heraud, J.M., Bessaud, M., 2022. Enterovirus detection in different regions of Madagascar reveals a higher abundance of enteroviruses of species C in areas where several outbreaks of vaccine-derived polioviruses occurred. BMC Infect Dis 22, 821.

Razafindratsimandresy, R., Joffret, M.L., Delpeyroux, F., Heraud, J.M., 2014. First full genome sequence of a human enterovirus a120, isolated in madagascar. Genome Announc 2.

Razafindratsimandresy, R., Raharinantoanina, J., Andrianiriana, E., Andrianjakatsilavo, A.E., Ratsimbazafy, A., Randrianasolo, L., Razanajatovo, N.H., Andriamandimby, S.F., Heraud, J.M., Lacoste, V., 2026. Molecular Characterization of Measles and Rubella Virus Strains from the 2018–2019 Epidemic in Madagascar. Pathogens 15, 514.

Razanajatovo, I.M., Andrianomiadana, L., Habib, A., Randrianarisoa, M.M., Razafimanjato, H., Rakotondrainipiana, M., Andriantsalama, P., Randriamparany, R., Andriamandimby, S.F., Vonaesch, P., Sansonetti, P.J., Lacoste, V., Randremanana, R.V., Collard, J.-M., Heraud, J.-M., Investigators, o.b.o.t.A., 2023. Factors Associated with Carriage of Enteropathogenic and Non-Enteropathogenic Viruses: A Reanalysis of Matched Case-Control Data from the AFRIBIOTA Site in Antananarivo, Madagascar. Pathogens 12, 1009.

Razanajatovo, N.H., Guillebaud, J., Harimanana, A., Rajatonirina, S., Ratsima, E.H., Andrianirina, Z.Z., Rakotoariniaina, H., Andriatahina, T., Orelle, A., Ratovoson, R., 2018. Epidemiology of severe acute respiratory infections from hospital-based surveillance in Madagascar, November 2010 to July 2013. PLoS one 13, e0205124.

Razanajatovo, N.H., Randriambolamanantsoa, T.H., Rabarison, J.H., Randrianasolo, L., Ankasitrahana, M.F., Ratsimbazafy, A., Raherinandrasana, A.H., Razafimanjato, H., Raharinosy, V., Andriamandimby, S.F., 2022. Epidemiological patterns of seasonal respiratory viruses during the COVID-19 pandemic in Madagascar, March 2020–May 2022. Viruses 15, 12.

Razanajatovo, N.H., Randriambolamanantsoa, T.H., Randrianasolo, L., Rabarison, J.H., Raveloson, S.U., Barasaona, B., Ranoelison, N.N., Ratsimbazafy, A., Razafimanjato, H., Raveloharivony, F.A., 2025. Comparative Analysis of the Evolutionary Dynamics of Seasonal Influenza Viruses in Madagascar Before and Since the Pandemic Period of COVID-19. Influenza and Other Respiratory Viruses 19, e70110.

Razanajatovo, N.H., Richard, V., Hoffmann, J., Reynes, J.-M., Razafitrimo, G.M., Randremanana, R.V., Heraud, J.-M., 2011. Viral etiology of influenza-like illnesses in Antananarivo, Madagascar, July 2008 to June 2009. PloS one 6, e17579.

Rice, A.L., Sacco, L., Hyder, A., Black, R.E., 2000. Malnutrition as an underlying cause of childhood deaths associated with infectious diseases in developing countries. Bulletin of the World Health organization 78, 1207–1221.

Rice, B.L., Golden, C.D., Randriamady, H.J., Rakotomalala, A.A.N.A., Vonona, M.A., Anjaranirina, E.J.G., Hazen, J., Castro, M.C., Metcalf, C.J.E., Hartl, D.L., 2021. Fine-scale variation in malaria prevalence across ecological regions in Madagascar: a cross-sectional study. BMC public health 21, 1018.

Richter, J., Tryfonos, C., Panagiotou, C., Nikolaou, E., Koliou, M., Christodoulou, C., 2013. Newly emerging C group enteroviruses may elude diagnosis due to a divergent 5′-UTR. International Journal of Infectious Diseases 17, e1245–e1248.

Roux, S., Adriaenssens, E.M., Dutilh, B.E., Koonin, E.V., Kropinski, A.M., Krupovic, M., Kuhn, J.H., Lavigne, R., Brister, J.R., Varsani, A., Amid, C., Aziz, R.K., Bordenstein, S.R., Bork, P., Breitbart, M., Cochrane, G.R., Daly, R.A., Desnues, C., Duhaime, M.B., Emerson, J.B., Enault, F., Fuhrman, J.A., Hingamp, P., Hugenholtz, P., Hurwitz, B.L., Ivanova, N.N., Labonté, J.M., Lee, K.-B., Malmstrom, R.R., Martinez-Garcia, M., Mizrachi, I.K., Ogata, H., Páez-Espino, D., Petit, M.-A., Putonti, C., Rattei, T., Reyes, A., Rodriguez-Valera, F., Rosario, K., Schriml, L., Schulz, F., Steward, G.F., Sullivan, M.B., Sunagawa, S., Suttle, C.A., Temperton, B., Tringe, S.G., Thurber, R.V., Webster, N.S., Whiteson, K.L., Wilhelm, S.W., Wommack, K.E., Woyke, T., Wrighton, K.C., Yilmaz, P., Yoshida, T., Young, M.J., Yutin, N., Allen, L.Z., Kyrpides, N.C., Eloe-Fadrosh, E.A., 2019. Minimum Information about an Uncultivated Virus Genome (MIUViG). Nature Biotechnology 37, 29–37.

Sauvage, V., Gomez, J., Boizeau, L., Vandenbogaert, M., Barbier, L., Tagny, C.T., Rakoto Alson, A.O., Bizimana, P., Coulibaly, S.O., Boullahi, M.A., Soumana, H., Mbensa, G., Caro, V., Laperche, S., 2021. New insights into Human Pegivirus-1 (HPgV-1) genotypes diversity in sub-Saharan Africa. Infect Genet Evol 94, 104995.

Schneider-Crease, I.A., Chiou, K.L., Moya, I.L., Lemma, A., Ayele, F., Haile, A.A., Kebede, F., Abebe, B., Bergman, T.J., Beehner, J.C., 2022. A novel statovirus identified in fecal samples from wild geladas in the Ethiopian highlands. Archives of virology 167, 2709–2713.

Seheri, L.M., Magagula, N.B., Peenze, I., Rakau, K., Ndadza, A., Mwenda, J.M., Weldegebriel, G., Steele, A.D., Mphahlele, M.J., 2018. Rotavirus strain diversity in Eastern and Southern African countries before and after vaccine introduction. Vaccine 36, 7222–7230.

Shu, Y., McCauley, J., 2017. GISAID: Global initiative on sharing all influenza data – from vision to reality. Eurosurveillance 22, 30494.

Song, K., Ye, R.-Z., Li, Y.-Y., Gong, C., Xia, L.-Y., Gao, W.-Y., Wang, Z.-F., Wang, N., Cheng, N., Zhang, M.-Z., Du, Y.-D., Xie, H., Zhu, D.-Y., Shi, W., Li, M.-Z., Zhao, L., Jia, N., Jiang, J.-F., Cui, X.-M., Huang, F., Cao, W.-C., 2024. A Statovirus-like virus from respiratory tracts of patients, China. Journal of Medical Virology 96, e29711.

Stewart, C.P., Fernald, L.C., Weber, A.M., Arnold, C., Galasso, E., 2020. Lipid-based nutrient supplementation reduces child anemia and increases micronutrient status in Madagascar: a multiarm cluster-randomized controlled trial. The Journal of nutrition 150, 958–966.

Stöver, B.C., Müller, K.F., 2010. TreeGraph 2: Combining and visualizing evidence from different phylogenetic analyses. BMC Bioinformatics 11, 7.

Taylor, L.J., Dotharda, M.I., Rubelb, M.A., Allena, A.A., Hwanga, Y., Rochea, A.M., Graham-Wootene, J., Fitzgeralde, A.S., Khatibe, L.A., Ranciaro, A., Thompson, S.R., Beggs, W.R., Campbell, M.C., Mokonef, G.G., Mpolokag, S.W., Fokunangh, C., Njamnshii, A.K., Mbunwec, E., Woldemeskelk, D., Belayk, G., Nyambol, T., Tishkoff, S.A., Collman, R.G., Bushman, F.D., 2021. Redondovirus Diversity and Evolution on Global, Individual, and Molecular Scales. Journal of Virology 95, e00817–00821.

Thieulent, C.J., Laverack, M., Carossino, M., Cronk, B., Caserta, L.C., Diel, D.G., Balasuriya, U.B.R., 2025. Detection of pathogens in dogs with respiratory disease during winter 2023–2024 using multiplex qPCR/RT-qPCR assays and next-generation sequencing. Frontiers in Veterinary Science Volume 12–2025.

Thom, K., Petrik, J., 2007. Progression towards AIDS leads to increased torque teno virus and torque teno minivirus titers in tissues of HIV infected individuals. Journal of Medical Virology 79, 1–7.

Timmerman, A.L., Schönert, A.L.M., Deijs, M., van Rijswijk, J., van Gils, M.J., van Goudoever, J.B., van Keulen, B.J., van der Hoek, L., 2025. Anellovirus constraint from 2 to 6 months postpartum followed by betatorquevirus and gammatorquevirus dominance in serum and milk. J Virol 99, e0084625.

Tirosh, O., Conlan, S., Deming, C., Lee-Lin, S.Q., Huang, X., Su, H.C., Freeman, A.F., Segre, J.A., Kong, H.H., 2018. Expanded skin virome in DOCK8-deficient patients. Nat Med 24, 1815–1821.

Tisza, M.J., Belford, A.K., Domínguez-Huerta, G., Bolduc, B., Buck, C.B., 2021. Cenote-Taker 2 democratizes virus discovery and sequence annotation. Virus Evolution 7.

Tisza, M.J., Petrosino, J.F., Cregeen, S.J.J., 2025. Cenote-Taker 3 for Fast and Accurate Virus Discovery and Annotation of the Virome. bioRxiv, 2025.2008. 2020.671380.

Tisza, M.J., Varsani, A., Petrosino, J.F., Cregeen, S.J.J., 2026. Cenote-Taker 3 for fast and accurate virus discovery and annotation of the virome. Peer Community Journal 6.

Tochetto, C., Cibulski, S.P., Muterle Varela, A.P., Cerva, C., Alves De Lima, D., Fumaco Teixeira, T., Quoos Mayer, F., Roehe, P.M., 2021. A variety of highly divergent eukaryotic ssDNA viruses in sera of pigs. Journal of General Virology 102.

Todd, A., Taylor, S., Huang, Q.S., 2015. Identification of Enterovirus C105 for the first time in New Zealand. Western Pacific surveillance and response journal: WPSAR 6, 60.

Tokarz, R., Hirschberg, D.L., Sameroff, S., Haq, S., Luna, G., Bennett, A.J., Silva, M., Leguia, M., Kasper, M., Bausch, D.G., 2013. Genomic analysis of two novel human enterovirus C genotypes found in respiratory samples from Peru. Journal of General Virology 94, 120–127.

Usman, M., Rana, M.S., Salman, M., Alam, M.M., Khurshid, A., Umair, M., Ullah, N., Anas, M., Ahmed, M., Ayub, A., 2024. Molecular characterization of rotavirus indicates predominance of G9P [4] genotype among children with acute gastroenteritis: First report after vaccine introduction in Pakistan. Journal of Medical Virology 96, e29761.

Van Leer-Buter, C.C., Poelman, R., Borger, R., Niesters, H.G., 2016. Newly Identified Enterovirus C Genotypes, Identified in the Netherlands through Routine Sequencing of All Enteroviruses Detected in Clinical Materials from 2008 to 2015. J Clin Microbiol 54, 2306–2314.

Varsani, A., Opriessnig, T., Celer, V., Maggi, F., Okamoto, H., Blomström, A.-L., Cadar, D., Harrach, B., Biagini, P., Kraberger, S., 2021. Taxonomic update for mammalian anelloviruses (family Anelloviridae). Archives of Virology 166, 2943–2953.

Volle, R., Razafindratsimandresy, R., Joffret, M.-L., Bessaud, M., Rabemanantsoa, S., Andriamamonjy, S., Raharinantoanina, J., Blondel, B., Heraud, J.-M., Bailly, J.-L., Delpeyroux, F., 2019. High Permissiveness for Genetic Exchanges between Enteroviruses of Species A, including Enterovirus 71, Favors Evolution through Intertypic Recombination in Madagascar. Journal of Virology 93, 10.1128/jvi.01667-01618.

Xiang, Z., Tsatsral, S., Liu, C., Li, L., Ren, L., Xiao, Y., Xie, Z., Zhou, H., Vernet, G., Nymadawa, P., Shen, K., Wang, J., 2014. Respiratory infection with enterovirus genotype C117, China and Mongolia. Emerg Infect Dis 20, 1075–1077.

Xiao, J., Wang, J., Zhang, Y., Sun, D., Lu, H., Han, Z., Song, Y., Yan, D., Zhu, S., Pei, Y., 2021. Coxsackievirus B4: an underestimated pathogen associated with a hand, foot, and mouth disease outbreak. Archives of virology 166, 2225–2234.

Yao, X., Zou, X., Cao, B., 2025. The Human Respiratory Virome in Health and Disease: Interactions, Dysbiosis, and Methodological Challenges. Advanced Genetics 6, e00022.

Ye, R.-Z., Zhao, J.-Q., Xie, H., Zhao, L., Gong, C., Wang, Z.-F., Yue, N., Xia, L.-Y., Song, K., Dong, B., 2025. A tombus-like virus in patients with lower respiratory tract infection: an observational study based on meta-transcriptomic sequencing. Emerging Microbes & Infections 14, 2494704.

Yolshin, N.D., Ivanova, A.A., Perederiy, A.A., Amosova, I.V., Timoshicheva, T.A., Stolyarov, K.A., Danilenko, D.M., Lioznov, D.A., Komissarov, A.B., 2026. Nationwide Genomic Surveillance of Human Respiratory Adenoviruses in 2023–2024: Evidence of Extensive Diversity and Recombination in Russia. Viruses 18, 136.

Zell, R., Delwart, E., Gorbalenya, A., Hovi, T., King, A., Knowles, N., Lindberg, A.M., Pallansch, M., Palmenberg, A., Reuter, G., 2017. ICTV virus taxonomy profile: Picornaviridae. Journal of General Virology 98, 2421–2422.

Zhang, M., Fan, S., Liang, M., Wu, R., Tian, J., Xian, J., Zhou, X., Chen, Q., 2024. A panoramic view of the molecular epidemiology, evolution, and cross-species transmission of rosaviruses. Vet Res 55, 145.

Zhu, Y., Sun, Y., Li, C., Lu, G., Jin, R., Xu, B., Shang, Y., Ai, J., Wang, R., Duan, Y., Chen, X., Xie, Z., 2024. Genetic characteristics of human parainfluenza viruses 1–4 associated with acute lower respiratory tract infection in Chinese children, during 2015–2021. Microbiology Spectrum 12, e03432–03423.

Zhuo, R., Parsons, B.D., Lee, B.E., Drews, S.J., Chui, L., Louie, M., Crago, B., Freedman, S.B., Ali, S., Pang, X., 2018. Identification of enteric viruses in oral swabs from children with acute gastroenteritis. The Journal of Molecular Diagnostics 20, 56–62.

Zohdy, S., Grossman, M.K., Fried, I.R., Rasambainarivo, F.T., Wright, P.C., Gillespie, T.R., 2015. Diversity and Prevalence of Diarrhea-Associated Viruses in the Lemur Community and Associated Human Population of Ranomafana National Park, Madagascar. International Journal of Primatology 36, 143–153.

